# Keratinocyte VISTA attenuates UV light-induced skin injury by suppressing cutaneous type I interferon (IFN-I) response

**DOI:** 10.64898/2025.12.04.692370

**Authors:** Zachary T. Peters, Lindsay K. Mendyka, J’Voughnn Blake, Himanshu B. Goswami, Angelique N. Cortez, Grace E. Crossland, Sicong Shan, Elizabeth C. Nowak, Sarkar Mrinal, E. Gudjodsson Johann, Christopher M. Burns, Dorothea T. Barton, Bruce R. Blazar, Tyler J. Curiel, Rodwell Mabaera, Victoria P. Werth, Andrea Kalus, Keith Elkon, Randolph J. Noelle, Sladjana Skopelja-Gardner

## Abstract

Persistent production of type I interferons (IFN-Is) is a hallmark of cutaneous lupus erythematosus (CLE). Ultraviolet (UV) light stimulates IFN-I response in the skin and exacerbates CLE. Here, we identify V-type immunoglobulin domain-containing suppressor of T cell activation (VISTA) as a negative regulator of both basal and UV-induced IFN-I responses in the skin and show that VISTA limits skin photosensitivity in an IFN-I-dependent manner, in part through Stimulator of Interferon Genes (STING). Furthermore, we demonstrate a novel role for VISTA in keratinocytes both at steady state and in response to UV light. Conditional deletion of VISTA in epidermal keratinocytes results in >10-fold increase in the basal skin IFN-I score and a heightened UV-induced skin injury score, which is dependent on IFN-I signaling. VISTA-targeting monoclonal antibodies suppress UV-induced IFN-I response in human keratinocytes and in mice expressing human VISTA *in vivo*, which also reduces UV-induced skin injury score. Our findings highlight VISTA as a potential therapeutic target for suppressing IFN-I responses in cutaneous lupus patients who suffer from UV-induced skin inflammation.

## Introduction

A defining molecular feature of autoimmune and inflammatory diseases such as systemic lupus erythematosus (SLE), cutaneous LE (CLE), and dermatomyositis is a persistently high type I interferon (IFN-I) signature in the skin (*1–4*). In SLE, the high skin IFN-I signature is detectable before the development of lesions, systemic IFN-I, or disease manifestations in other organs (*5, 6*). Sensitivity to ultraviolet (UV) rays from the sun, a condition known as photosensitivity, affects ∼70-80% of SLE and CLE patients and can exacerbate both local and systemic disease, including nephritis (*7–12*). Importantly, UV light is a potent trigger of IFN-I (*13*), and the UV-associated IFN-I gene signature is prominent even in non-lesional SLE skin (*14*). While blocking IFN-I signaling has some effects in CLE (*15*), no targeted therapies have been developed for photosensitivity.

Aberrant sensing of nucleic acids and defective clearance of apoptotic cells have been implicated in IFN-I production in lupus skin (*16–19*). Genetic variants in the regulators of IFN-I, such as STING, interferon regulatory factors 5 (IRF5) and 7 (IRF7), increase the risk of developing lupus and are associated with elevated IFN-I (*20–23*). In response to UV, apoptotic epidermal keratinocytes release cytoplasmic and mitochondrial DNA, stimulating IFN-I production via cGAS-STING, which further amplifies inflammation in lupus skin (*13, 24*). Epidermal keratinocytes are central to this process, as they both produce and respond to IFN-Is, and keratinocyte-derived IFN-κ has emerged as a key contributor to photosensitivity in CLE (*21, 25, 26*). Despite advances in defining the drivers of the IFN-I pathways, mechanisms that negatively regulate IFN-I production, particularly in keratinocytes and photosensitive SLE skin, remain poorly understood.

VISTA (V-type immunoglobulin domain-containing suppressor of T cell activation, also known as PD-1H, DD1α, or *VSIR*) is an immunoregulatory member of the B7 family of ligands and receptors, and its inhibitory functions have primarily been established in myeloid cells and T cells (*27–29*). In T cells, VISTA acts as a receptor, maintaining naïve T cell quiescence (*30*), reducing activation and production of IFNγ and IL2 (*27*), and enhancing differentiation of naïve T cells into regulatory T cells (*31*). In myeloid cells, VISTA promotes tolerogenic responses in macrophages (*32, 33*) and neutrophils (*34, 35*). While VISTA is also expressed by several stromal cell types, its immunosuppressive effects have primarily been studied in the context of cancer, where it is overexpressed in melanoma, ovarian cancer tumor cells, and melanoma-associated fibroblasts (*36*). Given its roles in maintaining peripheral tolerance and restricting inflammation, VISTA may play a role in autoimmune and chronic inflammatory conditions (*37, 38*). Deficiency in or blockade of VISTA results in a lupus-like skin disease in BALB/c mice and accelerates systemic disease in SLE mouse models (*35, 37, 39*). These pathologies were accompanied by an increase in the IFN-I signature (*35, 37*), suggesting VISTA may modulate the IFN-I response in lupus. However, whether VISTA directly regulates epithelial immune programs independent of systemic autoimmunity has not been addressed. Although VISTA is also expressed by keratinocytes (*40, 41*), its role in regulating basal and UV-induced IFN-I in the skin is unknown.

In this study, we investigate the role of VISTA in regulating UV-elicited skin IFN-I response and injury. Using C57BL/6 mice lacking VISTA (*Vsir*-/-), we interrogate the effects of VISTA loss on the basal cutaneous IFN-I tone and UV-triggered inflammatory and injury responses in the skin. To assess the translational relevance of our findings, we analyze transcriptomic changes in VISTA expression and IFN-I signaling in healthy and lupus human skin before and after UV *in vivo*. We investigate the regulatory functions of VISTA in the skin epidermis with conditional *Vsir* deletion in keratinocytes (K14^Cre^*Vsir^fl/fl^*).

Using single-cell transcriptomics, we identify cellular sources of basal and UV-induced IFN-I signature in the absence of keratinocyte VISTA. Finally, we explore the therapeutic potential of targeting human VISTA *in vivo* and in human keratinocytes *in vitro*, under IFN-I–rich conditions. These studies identify VISTA as an epidermal checkpoint that restrains STING–driven IFN-I and limits UV-provoked skin injury, highlighting a potential novel strategy to mitigate photosensitivity in CLE.

## Results

### VISTA deficiency amplifies basal skin interferon signature through STING

Previous studies have shown that global VISTA deficiency in BALB/c mice leads to the development of autoantibodies and skin lesions at ∼6-12 months of age (*35*). However, the mechanism(s) by which VISTA deficiency contributed to the CLE-like skin pathology were not established. Given that non-lesional skin in SLE and CLE patients exhibits prominent inflammatory signatures, including the IFN-I signature (*42–44*), we first investigated whether VISTA deficiency alters basal IFN-I signaling in young C57BL/6 (B6) mice, which do not develop spontaneous skin lesions. RNA-seq analysis of whole skin from wild-type (WT, B6) and VISTA-deficient (*Vsir*-/-) 3-month-old mice revealed 1188 differentially expressed genes in *Vsir*-/- skin (**Figure 1A**, **Table S1**). Many of the upregulated genes were interferon-stimulated genes (ISGs), and gene set enrichment analysis (GSEA) revealed *"Response to IFN-β"* as the most enriched pathway in *Vsir-/-* (NES = 2.571, p < 2.2 × 10^-16^), compared to B6 skin (**Figure 1A-C**). Quantitative PCR analysis confirmed elevated expression of multiple ISGs in *Vsir*-/- skin, with the cumulative IFN-I score showing ∼10-fold increase in VISTA-deficient compared to B6 skin (**Figure 1D**). Because IFN-κ has been implicated in CLE pathogenesis (*25, 45*), we asked whether the elevated skin IFN-I signature in the absence of VISTA reflects increased IFN-κ production. Both immunoblot and ELISA demonstrated higher IFN-κ protein levels in *Vsir*-/- compared to B6 skin (**Figure S1A-B**). VISTA deficiency as a model of CLE is not yet well characterized, so we next sought to understand how the strength of the skin IFN signature in *Vsir-/-* skin compares to that in established murine models of CLE-like skin disease and in human non-lesional CLE skin (*42*). We performed gene set variation analysis (GSVA) of bulk RNA-seq data from our (B6 vs. *Vsir*-/- skin) and three publicly available datasets: female 8-12 week old MRL/lpr non-lesional mouse skin (GSE222573), contralateral non-lesional ear of imiquimod (TLR7 agonist)-treated B6 mice (IMQ, GSE255519), and non-lesional human discoid LE (DLE) skin (GSE227329, GSE52471), relative to the respective control tissues in each dataset (*42*). GSVA demonstrated robust enrichment of IFN-I-related genes in *Vsir-/-* vs. B6 skin, akin to the enrichment observed in the skin of other lupus mouse models, as well as in non-lesional human DLE skin (**Figure 1E**). Moreover, this analysis revealed decreased expression of keratinocyte-associated genes and IL-17 signaling pathways in VISTA-deficient and human DLE, but not in MRL/lpr or IMQ-treated skin, highlighting additional similarities between VISTA-deficient and human CLE skin (**Figure 1E**).

**Figure 1.**
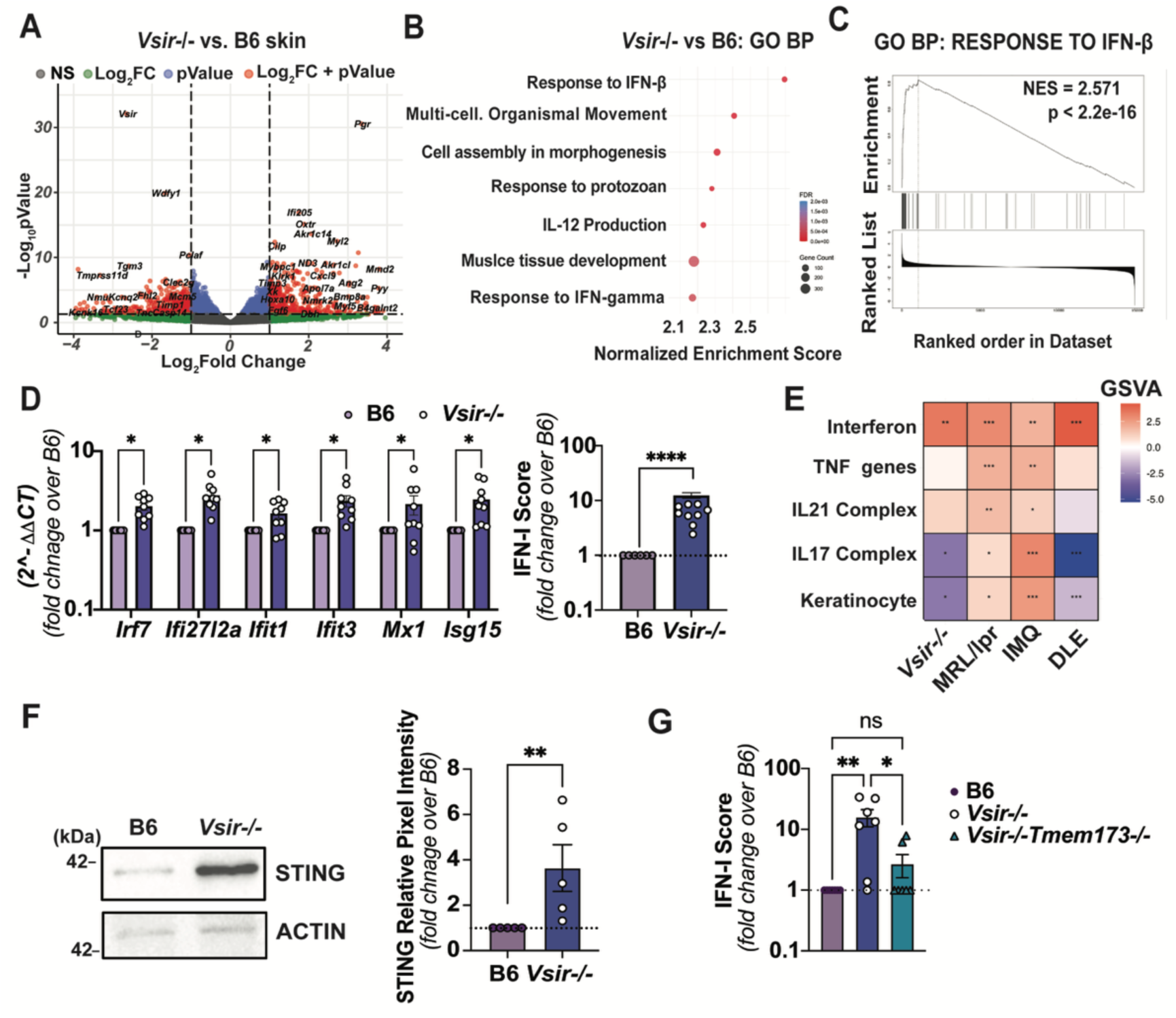
Heightened Type I interferon (IFN-I) signature in VISTA-deficient skin is in part mediated by STING. **(A)** Differentially expressed genes in VISTA-deficient (*Vsir*-/-), compared to wild-type (B6) murine skin from 12-week-old female mice; FDR-adjusted p-value < 0.01 and -1 < Log2FC > 1 (n = 3 per genotype). **(B)** Top differentially upregulated pathways from Gene set enrichment (GO) analysis: biological processes in *Vsir*-/- vs. B6 skin. Enrichment scores normalized to the size of the gene set, and p-values determined by permutation testing and multiple comparisons corrected for by Benjamini-Hochberg test (n=3). **(C)** Normalized enrichment score for GO biological pathway *Response to IFN*β in *Vsir*-/- vs. B6 skin. **(D)** Fold change in basal skin expression of individual IFN-I stimulated genes (ISGs, left) and cumulative IFN-I score (right), calculated as the sum of the normalized expression of 6 ISGs to β*-actin*, in *Vsir*-/-, relative to B6 skin. Statistical difference was determined by multiple Mann-Whitney U tests adjusted for false discovery rate using Benjamini, Krieger Yekutieli procedure (*p< 0.05, ****p<0.0001; n = 8) (**E)** Comparative gene set variation analysis (GSVA) showing hedges G effect size of pathways in *Vsir-/-*, MRL/lpr (GSE222573), contralateral ear in imiquimod (IMQ)- treated B6 mice (GSE255519), and human DLE skin (GSE227329, GSE52471) relative to B6, MRL/MpJ, untreated B6 ear, and human healthy control skin, respectively (*p<0.05, **p<0.01, ***p<0.001). (**F)** Representative immunoblot of basal STING protein in B6 and *Vsir-/-* whole skin protein extracts. Fold change in STING normalized to ACTIN was compared between genotypes by Mann-Whitney U test (**p< 0.01, n=5). (**G**) Differences in the basal skin IFN-I score, derived from genes in D, in B6, *Vsir*-/- and *Vsir-/-Tmem173*-/- mice (12 weeks, female), relative to B6 skin, were compared using One-way ANOVA with post-hoc Tukey test (n=7; *p< 0.05, **p<0.01, ns=not significant).

Activation of various nucleic acid-sensing pathways may underlie the increased IFN-I production in *Vsir*-/- skin (*46*). However, recent studies attributed the high IFN-I production in CLE skin to increased signaling through the Stimulator of Interferon Genes (STING), an adapter protein central to cytosolic DNA sensing (*20, 47*). Consistent with this, STING protein levels were increased in the skin of *Vsir*-/- mice compared to VISTA- sufficient B6 controls (**Figure 1F**). To directly assess the contribution of STING to the heightened IFN-I production in the absence of VISTA, we generated mice doubly deficient in VISTA and STING (*Vsir-/-Tmem173-/-*). Loss of STING reduced ISG expression and the cumulative IFN-I score in *Vsir-/- Tmem173*-/- mice by ∼80% relative to *Vsir-/-* controls (**Figure 1G and Figure S1C**). Taken together, these data identify VISTA as a suppressor of STING-mediated IFN-I production in the skin.

### VISTA regulates UV light-induced IFN-I responses and cellular inflammation

Exposure to sunlight can provoke and exacerbate disease in skin conditions marked by a persistently high IFN-I signature, such as CLE (*48, 49*). Acute exposure to UV light stimulates a potent IFN-I response in healthy human and murine skin *in vivo* (*13*). In healthy murine skin, the IFN-I response subsides over time, suggesting the presence of negative regulatory mechanisms (*13*). By contrast, non-lesional lupus skin exhibits a persistently high basal cutaneous IFN-I signature (*25*), and UV exposure further amplifies ISG expression in lupus compared to healthy skin (*13*). Given the potential suppressive role of VISTA in cutaneous IFN-I production (**Figure 1**), we asked whether VISTA regulates the magnitude of the IFN-I response to an acute dose of UV light delivered to the shaved dorsal section of B6 and *Vsir-/-* mice (**Figure 2A).** RNA-seq analysis revealed 381 differentially expressed genes in *Vsir-/-* compared to B6 skin 3 hours after UV (**Figure S2A** and **Table S1).** GSEA identified “*Response to IFN-β"* as the top upregulated pathway in *Vsir-/-* compared to B6 skin after UV (**Figure 2B**). Furthermore, GSVA showed greater upregulation of IFN-I and IFN-γ signatures in *Vsir*-/- skin, alongside suppression of keratinocyte-associated genes (**Figure S2B**). To confirm the heightened UV-induced IFN-I response in VISTA-deficient skin, we quantified the cumulative IFN-I score by qPCR. While exposure to UV stimulated ISG expression in both B6 and *Vsir-/-* skin, the IFN-I score was ∼5-fold higher in *Vsir*-/-, relative to B6 skin, 3 hours after UV (**Figure 2C, S2C)**.

**Figure 2.**
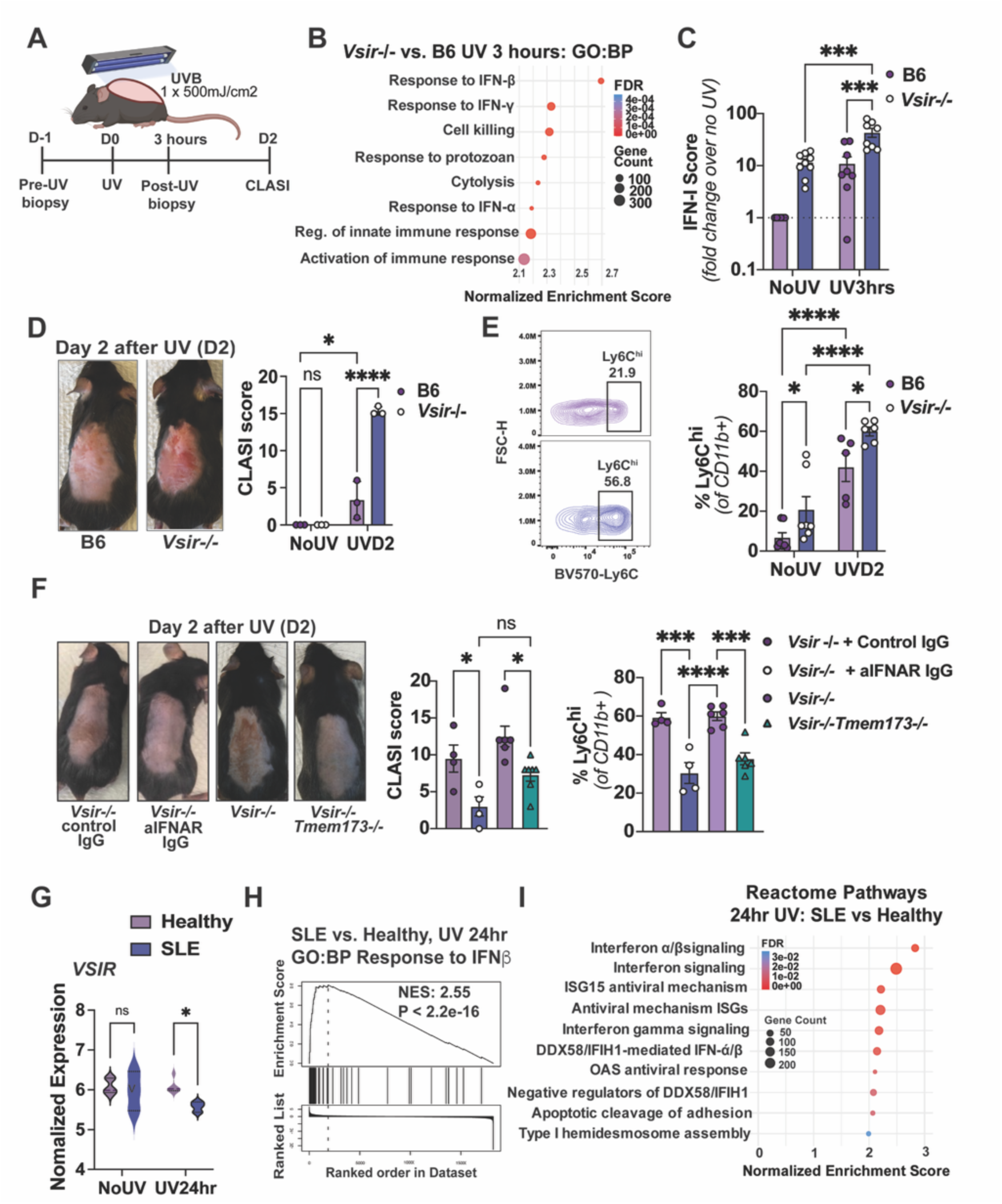
Loss of VISTA exacerbates UV-triggered skin injury through increased IFN-I activity. **(A)** Schematic of UVB exposure and skin collection. Shaved dorsal back of B6 and *Vsir*-/- mice (12 weeks, female) was exposed to one dose of UVB (500mJ/cm^2^). Skin was collected before (NoUV) and 3hr after (UV3hrs) UVB for analyses in B and C. CLASI scores and inflammatory monocyte infiltration into the skin were evaluated on day 2 (UVD2) after UVB in D-F. **(B)** Gene set enrichment analysis (GSEA) of the most highly upregulated GO: Biological processes pathways in *Vsir-/-* mouse skin 3hr after UV, compared to B6 skin. Enrichment scores are normalized to the size of the gene set, and p-values determined by permutation testing and multiple comparisons corrected for by Benjamini-Hochberg test (n=3). **(C)** Differences in the skin IFN-I score of B6 and *Vsir*-/- mice before (NoUV) and 3hr after UV, relative to unexposed B6 skin, were compared by two-way ANOVA with post-hoc Tukey test (n=8, ***p<0.001). IFN-I score was calculated as the sum of the normalized expression of 6 ISGs (*Irf7, Ifi27l2a, Ifit1, Ifit3, Isg15, Mx1*) to β*-actin*. **(D)** Representative images of mouse skin on day 2 after UV. **(D-E)** Differences in (D) Cutaneous Lupus Activity and Severity Index (CLASI) scores and (E) the percentage of Ly6C^hi^ inflammatory monocytes within myeloid cells (CD11b+) in B6 and *Vsir*-/- mice before (NoUV) and 2 days after UV (UVD2) were compared by two-way ANOVA (n = 3, *p<0.05, ****p<0.0001, ns=not significant). **(F)** Differences in the CLASI scores and percent Ly6C^hi^ inflammatory monocytes on day 2 after UV in *Vsir*-/- mice that received anti-IFNAR or control IgG 3hr post-UV, or *Vsir-/-* and *Vsir*-/-*Tmem173*-/- mice were compared by one-way ANOVA with post-hoc Tukey Test (n=4-6, *p<0.05, ***p< 0.001, ****p<0.0001, ns = not significant**) (G)** *VSIR* expression in healthy and SLE skin before (No UV) and 24 hours after acute UVB (2 x MED, two-way ANOVA with fisher’s LSD test; n=3, *p< 0.05). Top **(H)** GO: Biological Processes **(I)** and Reactome pathways enriched in the SLE patient skin 24 hr after UV (GSEA). Enrichment scores are normalized to size of gene set and p values determined by permutation testing and multiple comparisons corrected for by Benjamini-Hochberg test (n=3).

Given the previously proposed role of VISTA in the clearance of dead cells (*38*), we assessed UV-induced apoptosis by TUNEL staining. *Vsir-/-* and B6 skin had comparable prevalence of apoptotic cells at baseline, 3 hours, and 1 day following UV (**Figure S2D**). Therefore, these data suggest that VISTA deficiency permits an exaggerated UV-induced IFN-I independently of epidermal apoptosis.

To understand if the heightened IFN-I response in *Vsir-/-* mice predisposes the skin to increased UV-induced inflammation or injury, as seen in lupus patients, we monitored the mice for two additional days after UV and compared the extent of skin injury between genotypes. At this later time point, *Vsir-/-* mice demonstrated significantly higher Cutaneous Lupus Activity and Severity Index (CLASI) score (*50*), which accounts for redness, scaling, and scarring, compared to the VISTA-sufficient mice (**Figure 2D**). Moreover, *Vsir*-/- skin showed greater inflammation, reflected by an increased frequency of inflammatory Ly6C^hi^ monocytes (**Figure 2E, S2E**). To test whether the photosensitive skin response in the absence of VISTA was due to enhanced IFN-I signaling, we treated *Vsir-/-* mice with a blocking antibody to the IFN-I receptor alpha (anti-IFNAR IgG) or control IgG 3 hours after UV exposure, the time of peak early ISG expression. IFNAR blockade significantly reduced UV-induced CLASI scores and decreased the prevalence of Ly6C^hi^ inflammatory monocytes in *Vsir-/-* skin (**Figure 2F)**. Because STING mediated much of the basal IFN-I activity in *Vsir-/-* skin (**Figure 1G**), we next asked whether STING was required for the photosensitive response in the absence of VISTA. Compared to *Vsir-/-* mice, *Vsir-/-Tmem173-/-* mice exhibited a partial but significant reduction in CLASI scores and inflammatory monocytes two days after UV (**Figure 2F**). Together, these data identify VISTA as a regulator of UV-induced cutaneous IFN-I response and implicate STING-mediated IFN-I as a key driver of exaggerated photosensitive skin injury observed in the absence of VISTA.

To assess the relevance of VISTA to human lupus photosensitivity, we examined VISTA transcript (*VSIR)* levels and IFN-I pathway activation in healthy and SLE skin 24 hours after acute UV exposure. RNA-seq analysis was performed on paired skin biopsies from healthy and SLE volunteers before and 24 hours after UV (two minimal erythema doses, GSE148535 (*13*)). Interestingly, *VSIR* transcript levels were significantly lower in SLE compared to healthy skin 24 hours after UV (**Figure 2G**). This reduction in *VSIR* expression was accompanied by a significantly stronger IFN-I gene signature 24 hours after UV in SLE compared to healthy skin (**Figure 2H**). In fact, the top differentially upregulated pathways in SLE relative to healthy skin after UV were all related to IFN signaling (**Figure 2I**). Together, the findings in murine and human lupus skin suggest that VISTA suppresses cutaneous IFN-I production and that lower VISTA expression in UV-exposed SLE skin may contribute to the exaggerated IFN-I activation and photosensitive response.

### A keratinocyte-specific role for VISTA in the regulation of cutaneous IFN-I

Both hematopoietic and non-hematopoietic cells express VISTA (*41, 51–53*), including keratinocytes (*40, 41*). Previous studies have implicated keratinocytes in CLE pathogenesis and photosensitive responses, identifying them both as producers of and responders to IFN-Is, contributing to the heightened IFN-I signature in CLE skin (*4, 25*). Having confirmed VISTA expression in murine keratinocytes (**Figure S3A**), we first investigated whether VISTA plays a role in the regulation of basal IFN-I production in keratinocytes *in vitro*. Epidermal keratinocytes isolated from *Vsir-/-* tails and cultured for 5-7 days before protein isolation expressed more than 2-fold higher IFN-k levels compared to VISTA-sufficient (WT) keratinocytes (**Figure 3A)**. Although these *in vitro* findings suggest a keratinocyte-specific role for VISTA in regulating IFN-I production, they lack the tissue-specific context to capture the complex multicellular interactions in the skin. To address this *in vivo*, we generated mice with keratinocyte-specific VISTA deletion using the Cytokeratin 14 (K14) Cre-lox system (K14^Cre^*Vsir*^fl/fl^, (*54*)). Akin to the global VISTA deficiency, RNA-seq followed by GSVA identified “*Response to IFN-β”* as the top upregulated pathway in K14^Cre^*Vsir*^fl/fl^ skin relative to *Vsir*^fl/fl^ controls (**Figure 3B**). We confirmed that the basal skin IFN-I signature was increased in the absence of keratinocyte VISTA by qPCR analysis of ISGs, which revealed ∼10-fold higher IFN-I score in K14^Cre^*Vsir*^fl/fl^, compared to the cre-negative skin (**Figure 3C, Figure S3B)**. These findings indicated that keratinocyte-restricted VISTA loss is sufficient to drive the heightened skin IFN-I response seen with global VISTA deficiency.

**Figure 3.**
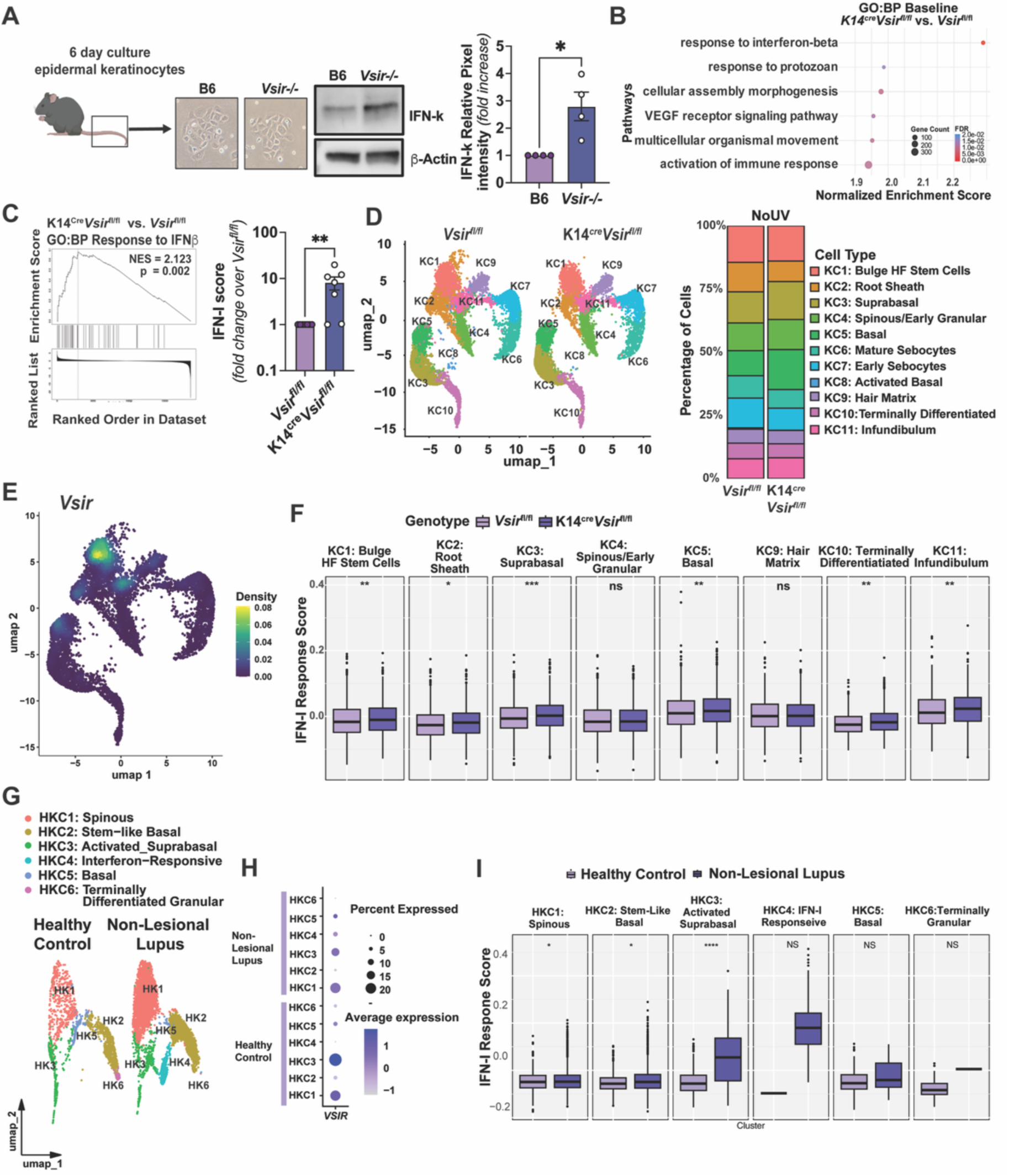
VISTA deficiency in keratinocytes results in a heightened skin IFN-I signature. **(A)** Schematic of isolation and culture of keratinocytes from B6 and *Vsir-/-* mouse tails. Baseline IFN-κ protein in B6 and *Vsir*-/- keratinocyte lysates after 6 days in culture, quantified relative to β-actin was compared by Mann-Whitney U Test (n=4, *p<0.05). **(B)** Top pathways enriched in K14^Cre^*Vsir*^fl/fl^ vs *Vsir^fl^*^/fl^ mouse skin by gene set enrichment analysis (GO: Biological Processes). Enrichment scores are normalized to size of gene set and p values determined by permutation testing and multiple comparisons corrected by Benjamini-Hochberg (n=4 mice per genotype). **(C)** Basal IFN-I scores in *Vsir^fl^*^/fl^ and K14^Cre^*Vsir*^fl/fl^ skin were compared by Mann-Whitney U Test (n= 7, **p<0.01). IFN-I score was calculated as the sum of the normalized expression of 6 ISGs (*Irf7, Ifi27l2a, Ifit1, Ifit3, Isg15, Mx1*) to β*-actin*. **(D)** Uniform manifold approximation and projection (UMAP) of keratinocyte clusters from *Vsir^fl^*^/fl^ and K14^Cre^*Vsir*^fl/fl^ skin scRNA-seq data generated with 10X FLEX platform. The distribution of keratinocyte clusters is shown in bar graphs. **(E)** Density plot of *Vsir* expression in keratinocyte clusters. **(F)** IFN-I response scores for keratinocyte clusters from *Vsir^fl^*^/fl^ and K14^Cre^*Vsir*^fl/fl^ skin were compared by multiple Wilcoxon tests corrected by Benjamini-Hochberg (n=4 mice per genotype; *p<0.05, **p<0.01, ***p<0.001, ns=not significant). (**G)** UMAP of healthy control and non- lesional (NL) lupus keratinocyte populations (scRNA-seq dataset GSE186476). (**H)** Dot plot of *VSIR* expression in healthy control and NL lupus patient skin keratinocyte clusters**. (I)** IFN-I response scores in Healthy control and NL lupus patient skin keratinocyte clusters were compared by multiple Wilcoxon Tests corrected by Benjamini-Hochberg (n=14 healthy control; n=7 NL Lupus; *p<0.05, ****p<0.0001, ns = not significant).

Since keratinocytes consist of different cellular subsets and many stromal and immune cells in the skin are responsive to IFN-Is, we next asked if the keratinocyte-specific function of VISTA is confined to a particular keratinocyte subset and which cells contribute to the elevated IFN-I signature in K14^Cre^*Vsir*^fl/fl^ skin. To address these questions, we performed single-cell RNA-seq analysis of K14^Cre^*Vsir*^fl/fl^ and *Vsir*^fl/fl^ skin, capturing eleven keratinocyte subsets: bulge hair follicle (HF) stem cells (KC1), root sheath (KC2), suprabasal (KC3), spinous/early granular (KC4), basal (KC5), mature and early sebocytes (KC6-7), activated basal (KC8), hair matrix (KC9), terminally differentiated (KC10), and infundibulum (KC11) (**Figure 3D, Figure S3C, and Table S1**). VISTA expression varied across keratinocyte subsets, with the highest *Vsir* transcript levels in follicular (KC1, KC2, KC9, KC11), and some expression in basal (KC5) cells (**Figure 3E**). Deletion of keratinocyte-specific VISTA altered subset composition, with a reduction in the proportion of follicular keratinocytes, the populations with the highest *Vsir* expression, in K14^cre^*Vsir*^fl/fl^ skin (**Figure 3D**). Analysis of the IFN-I response gene signature across keratinocyte subsets and genotypes revealed a significantly higher IFN-I response score in several keratinocyte clusters, including bulge HF stem cells (KC1), basal (KC5), and suprabasal (KC3) in K14^cre^*Vsir*^fl/fl^, compared to cre-negative (*Vsir^fl/fl^*) keratinocytes (**Figure 3F**). Beyond elevated IFN-I signaling, differential gene expression analysis showed that genes involved in the recruitment of myeloid cells (*Ccl9, Ccl24*), inflammation (*Il33, Il23, Tslp*), oxidative stress response (*Gsta2, Ch25h*), as well as antigen presentation (*H2-m2, H2- ob1, H2-aa, H2-q4*), were significantly upregulated in the keratinocytes from K14^Cre^*Vsir*^fl/fl^ skin (**Figure S3D, Table S1**). Pathway analysis revealed these transcriptional changes were associated with increased expression of leukocyte recruitment, adhesion, and antigen presentation pathways and decreased expression of keratinocyte and epidermal development pathways in K14^Cre^*Vsir*^fl/fl^ keratinocytes, compared to VISTA-sufficient controls (**Figure S3D-E**). Together, these data suggest that the absence of keratinocyte VISTA results in changes to keratinocyte subsets and increased basal IFN-I and inflammatory signaling in keratinocytes.

Because epidermal keratinocytes in non-lesional CLE skin exhibit a strong IFN-I signature (*4, 25*), we asked whether *VSIR* expression may be linked to the high IFN-I signature in CLE keratinocytes by analyzing a single-cell RNA-seq dataset of healthy and CLE skin (GSE186476, (*4*)). In human keratinocytes, *VSIR* was most highly expressed in spinous (HKC1) and suprabasal (HKC3) keratinocyte subsets (**Figure 3G-H, Figure S3F**). Very few follicular keratinocytes, high expressors of *Vsir* in murine skin, were captured, precluding the analysis of that subset. Notably, *VSIR* levels were reduced across the different *VSIR*-expressing keratinocyte subsets in non-lesional CLE, compared with healthy skin, particularly in the activated suprabasal cluster (**Figure 3H**). Lower *VSIR* levels in CLE keratinocytes were associated with increased transcriptional scores of IFN-I-related genes in the activated suprabasal keratinocyte subset, the highest *VSIR* expressor (**Figure 3I**). Thus, similarly to the findings in the K14^Cre^*Vsir*^fl/fl^ skin, reduced VISTA expression in keratinocytes is also associated with a heightened IFN-I activity in human CLE.

### Keratinocyte-expressed VISTA protects the skin from UV-induced, IFN-I-mediated injury

UVB light directly impacts the epidermis and stimulates IFN-I production by keratinocytes (*55*). Since keratinocyte-specific VISTA loss was sufficient to elevate the basal cutaneous IFN-I signature, we asked if keratinocyte-expressed VISTA is required to suppress UV-induced IFN-I response and skin injury, by comparing the skin response to UV light in K14^Cre^*Vsir*^fl/fl^ and *Vsir*^fl/fl^ (VISTA-sufficient) mice. Two days after UV, K14^Cre^*Vsir*^fl/fl^ mice developed ∼2-fold higher CLASI scores than the *Vsir*^fl/fl^ controls, comparable in magnitude to those seen in the mice with global VISTA deficiency (**Figure 4A**). To examine how keratinocyte VISTA influences stromal and immune responses to UV light, we performed single-cell RNA-seq on K14^Cre^*Vsir*^fl/fl^ and *Vsir*^fl/fl^ skin collected 24 hours after UV (**Figure 4B**). Keratinocyte subset analysis revealed increased prevalence of basal keratinocytes, including the activated basal subset, and a decrease in the proportion of follicular keratinocyte subsets (bulge and root sheath) in K14^Cre^*Vsir*^fl/fl^, compared to *Vsir*^fl/fl^ skin (**Figure 4B**). In VISTA-sufficient skin, UV exposure upregulated *Vsir* expression in several keratinocyte subsets, particularly bulge HF stem cells (KC1) and basal (KC5) keratinocytes (**Figure 4C**). In the absence of VISTA, bulge HF stem cells (KC1) and activated basal keratinocytes (KC8) exhibited significantly higher IFN-I response scores in K14^Cre^*Vsir*^fl/fl^ compared to *Vsir*^fl/fl^ skin (**Figure 4D**). Unbiased gene expression and pathway analysis identified *Response to IFN-β* as a top upregulated pathway in VISTA-deficient keratinocytes after UV (**Figure 4E-F**), alongside granulocyte and leukocyte chemoattractant programs (**Figure 4F**). In contrast, wound healing pathways, including *extracellular matrix organization* and *collagen formation*, as well as keratinocyte differentiation pathways, such as *hair cycle and basement membrane organization,* were suppressed in K14^Cre^*Vsir*^fl/fl^ keratinocytes, compared to *Vsir*^fl/fl^ controls (**Figure 4F**). Together, these results suggest that keratinocyte VISTA not only limits UV-induced IFN-I responses but also may support keratinocyte repair programs in response to UV.

**Figure 4.**
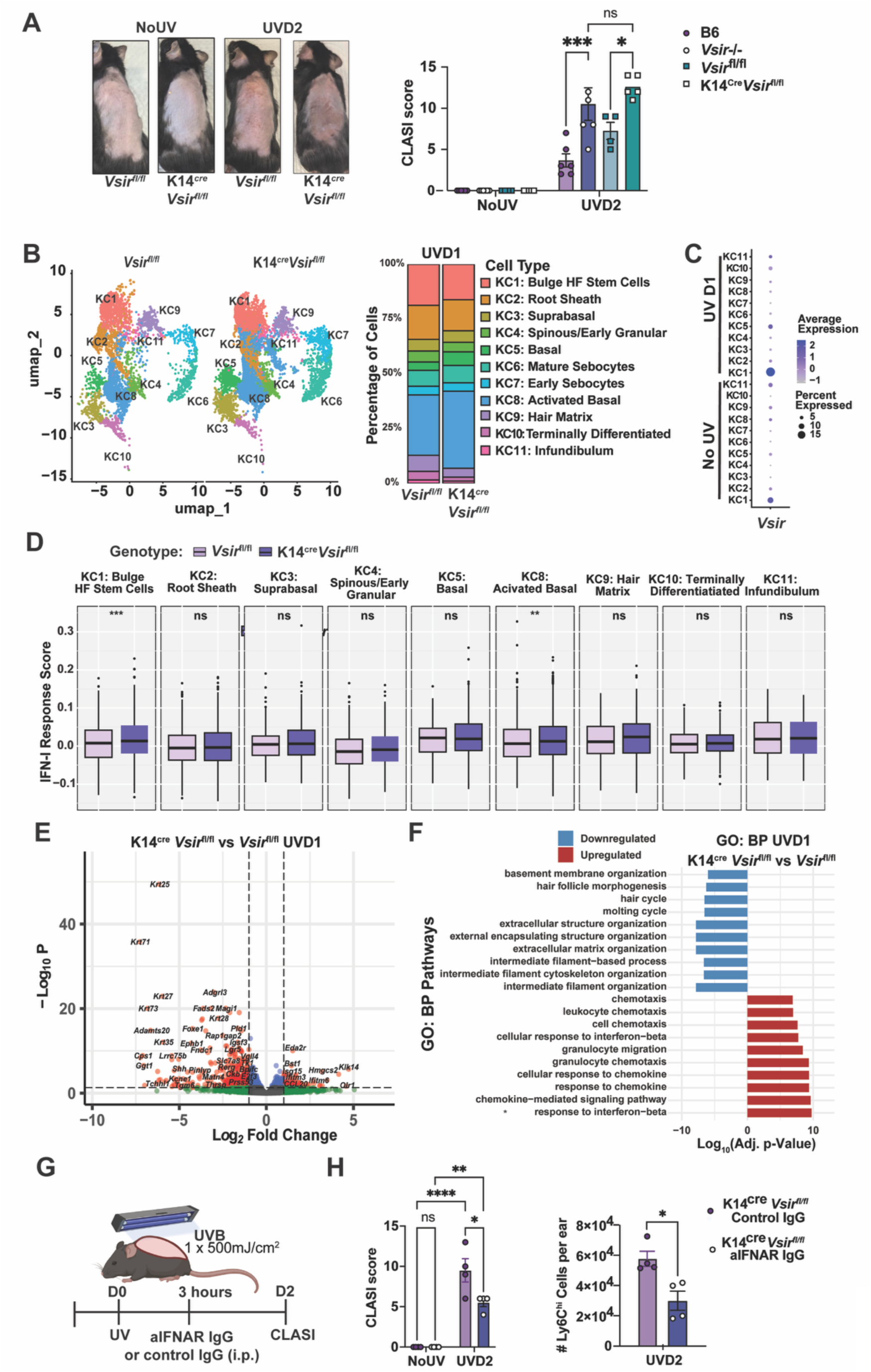
Keratinocyte-specific VISTA deficiency is sufficient to drive IFN-I– dependent exacerbation of UV-induced skin damage. **(A)** Representative images and CLASI scores of K14^Cre^*Vsir^fl^*^/fl^ and *Vsir^fl^*^/fl^ mice before and 2 days post-UV (1 x 500mJ/cm^2^) compared by two-way ANOVA with post-hoc Tukey test (n=5, *p<0.05, ***p<0.001, ns = not significant). **(B)** UMAP of K14^Cre^*Vsir*^fl/fl^ and *Vsir*^fl/fl^ mouse keratinocyte clusters from the skin collected on day 1 after UV, split by genotype; relative distribution of each cluster within keratinocytes is represented in the bar graphs. **(C)** Dot plot of *Vsir* expression in each keratinocyte cluster from *Vsir*^fl/fl^ unexposed skin (No UV) or skin collected one day after UV (UVD1, 250mJ/cm^2^). **(D)** IFN-I response gene signature scores in keratinocyte clusters from K14^Cre^*Vsir^fl^*^/fl^ and *Vsir^fl^*^/fl^ skin were compared by multiple paired cluster-wise Wilcoxon tests corrected by Benjamini-Hochberg (n=4 mice per genotype; **p<0.01,***p<0.001, ns = not significant). **(E)** Differentially expressed genes in K14^Cre^*Vsir^fl^*^/fl^ vs. *Vsir^fl^*^/fl^ keratinocyte clusters one day after UV (FDR adjusted p value < 0.01 and -1 < Log2FC > 1). **(F)** Gene set enrichment analysis (GSEA) of the top 10 GO: Biological processes pathways in K14^Cre^*Vsir^fl^*^/fl^ mouse keratinocyte cluster on day 1 after UV, compared to *Vsir*^fl/fl^ keratinocytes. Enrichment scores are normalized to size of gene set and p values determined by permutation testing and corrected for multiple comparisons by Benjamini-Hochberg (n=4). **(G)** Schematic of IFNAR blockade following UV. Shaved dorsal back of K14^Cre^*Vsir*^fl/fl^ mice (12 weeks, female) were exposed to one dose of UVB (500mJ/cm^2^) and anti-IFNAR (aIFNAR IgG) or control IgG administered i.p. (200µg) 3 hours later. **(H)** CLASI scores of K14^Cre^*Vsir*^fl/fl^ mice that received IFNAR blocking antibody were compared to those of mice that received control IgG by two-way ANOVA with post-hoc Tukey test (n=4, *p<0.05, **p<0.01, ****p<0.0001, ns = not significant). The number of Ly6C^hi^ monocytes in the skin of K14^Cre^*Vsir*^fl/fl^ mice that received IFNAR blocking antibody was compared to that of mice that received control IgG by unpaired t-test (n=4, *p<0.05).

Finally, to test whether the photosensitivity observed in K14^Cre^*Vsir*^fl/fl^ skin is due to the heightened IFN-I signaling, we treated K14^Cre^*Vsir*^fl/fl^ mice with anti-IFNAR or control IgG 3 hours after exposure to UV (**Figure 4G**). As seen in global VISTA deficiency (*Vsir-/-*, **Figure 2F**), IFNAR blockade significantly reduced CLASI scores and decreased the infiltration of inflammatory Ly6C^hi^ monocytes in K14^Cre^*Vsir*^fl/fl^ skin (**Figure 4H**). Therefore, these data suggest that keratinocyte-intrinsic VISTA constrains UV-induced IFN-I response and protects against inflammatory skin injury, while also promoting keratinocyte integrity and wound-healing responses.

### Keratinocyte VISTA deficiency reshapes stromal and immune cell phenotypes and disrupts cell–cell communication after UV

Single-cell RNA-seq analysis of K14^Cre^*Vsir*^fl/f^ and *Vsir*^fl/f^ skin identified 11 cell types in addition to keratinocytes, including fibroblasts, myocytes, melanocytes, Langerhans cells, macrophages, T cells, neutrophils, endothelial cells, adipocytes, and mast cells/basophils (**Figure 5A and Figure S4A**). At baseline, we observed decreased keratinocytes and melanocytes, but increased frequency of T cells and fibroblasts in K14^Cre^*Vsir*^fl/f^ vs *Vsir*^fl/f^ skin (**Figure S4B**). One day after UV, K14^Cre^*Vsir*^fl/fl^ skin had an increased proportion of monocytes and fibroblasts and a decreased proportion of melanocytes, compared to control *Vsir*^fl/fl^ skin (**Figure 5A**). To understand which cells contributed to the heightened IFN-I signature in the absence of keratinocyte-expressed VISTA, we compared the strength of the IFN-I response gene score across clusters and genotypes (**Figure 5B**). In addition to the keratinocytes, fibroblasts, monocytes, melanocytes, macrophages, adipocytes, and mast/basophils had increased IFN-I response gene scores in K14^Cre^*Vsir*^fl/fl^ skin, compared to their counterparts in VISTA- sufficient skin (**Figure 5B**). An unbiased analysis of the differentially expressed pathways revealed that IFN-I-related pathways were among the most upregulated in fibroblasts and macrophages after UV, in the absence of keratinocyte-expressed VISTA (**Figure 5C-D**). Moreover, pathways of *chemokine production* and *chemotaxis* were enriched in fibroblasts and monocytes from K14^Cre^*Vsir*^fl/fl^ skin, respectively (**Figure 5C-E**). To determine if the transcriptional changes in fibroblasts, macrophages, and monocytes resulting from VISTA loss in keratinocytes affect the phenotypic composition of these populations, we compared the distribution of different cellular subtypes within each cluster across genotypes (**Figure 5C-E, S4C-E**). Fibroblasts in K14^Cre^*Vsir*^fl/fl^ skin showed increased prevalence of secretory and inflammatory clusters, and reduced hair follicle-associated fibroblasts (**Figure 5C**). Interferon-activated and inflammatory monocyte subsets were more prevalent in K14^Cre^*Vsir*^fl/fl^ skin (**Figure 5E**). Therefore, in addition to its keratinocyte-intrinsic effects (**Figures 3 and 4**), the loss of keratinocyte VISTA enhanced IFN-I and inflammatory responses to UV in multiple stromal and immune populations in the skin, as well as altered their phenotypes and composition.

**Figure 5.**
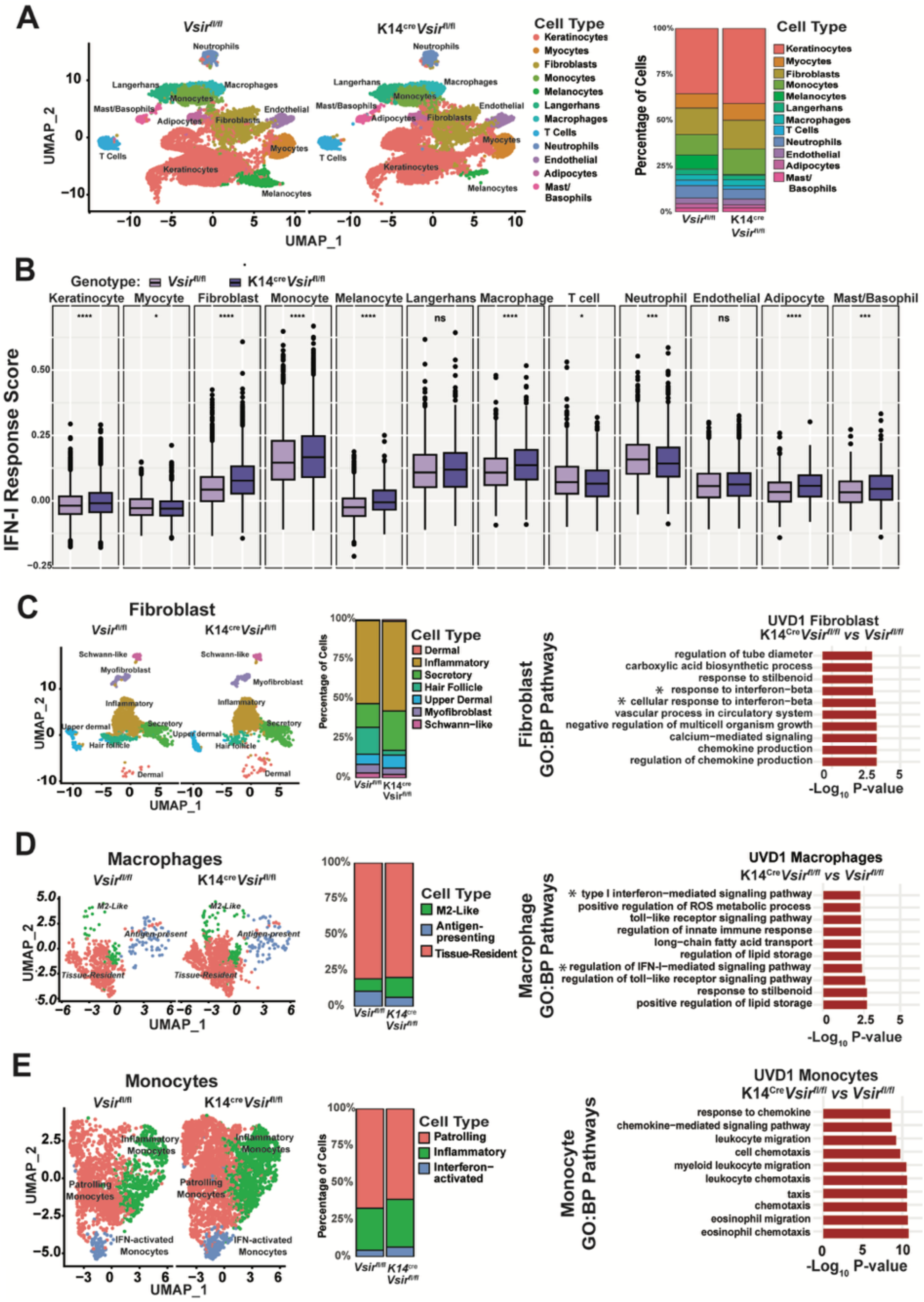
Loss of keratinocyte VISTA results in elevated IFN-I response across skin cell types and enhances UV-induced monocyte recruitment. **(A)** UMAP of K14^Cre^*Vsir*^fl/fl^ and *Vsir*^fl/fl^ skin cell clusters on day 1 after UVB (1 x 250mJ/cm^2^), split by genotype; relative percentage of each cluster is represented in the bar graphs. **(B)** IFN-I response gene signature scores in clusters from K14^Cre^*Vsir^fl^*^/fl^ and *Vsir^fl^*^/fl^ skin were compared by multiple paired cluster-wise Wilcoxon tests with Benjamini-Hochberg (n=4 mice per genotype; *p<0.05, ***p<0.001, ****p<0.0001, ns = not significant). **(C-E)** UMAP visualization of (C) fibroblast, (D) macrophage, and (E) monocyte clusters in K14^Cre^*Vsir*^fl/fl^ vs. *Vsir*^fl/fl^ skin. Bar graphs illustrate the relative proportion of each subcluster within the main population. Enriched GO biological process pathways in K14^Cre^*Vsir*^fl/fl^ vs. *Vsir*^fl/fl^ cells are displayed for each cluster. Enrichment scores were normalized to gene set size, and p-values were calculated by permutation testing with Benjamini-Hochberg (n = 4 mice per genotype).

Ligand–receptor interaction analysis showed that keratinocyte-specific VISTA deficiency disrupted intercellular communication within the skin. Broadly, the number and strength of imputed cell-cell contact interactions were reduced across cell types in K14^Cre^*Vsir*^fl/fl^, compared to *Vsir*^fl/fl^, skin one day after UV (**Figure S5A-B**). However, secreted signaling was overall increased in K14^Cre^*Vsir*^fl/fl^ skin, particularly between recruited cell types, such as monocytes and neutrophils, and skin-resident cells (**Figure S5C-D**). Many pathways associated with wound healing, keratinocyte migration, and immune regulation were decreased in K14^Cre^*Vsir*^fl/fl^ skin after UV (e.g. Thy1 and NOTCH; Alcam, Cadherins (CDH), APP and JAM (*56–59*); ICAM, PECAM, ESAM, SELPG, **Figure S5B**). Interestingly, decreased *Jam* signaling was attributed to diminished interactions between keratinocytes, melanocytes, and immune cells (**Figure S5E**). In parallel, secreted *Angptl* signaling was increased overall in K14^Cre^*Vsir*^fl/fl^ skin (**Figure S5F**). Taken together, these results implicate keratinocyte VISTA in regulating skin responses to UV light by suppressing IFN-I activity, preserving homeostatic cell–cell interactions, and limiting proinflammatory crosstalk between stromal and immune cells.

### Targeting human VISTA suppresses UV-induced IFN-I responses in the skin and keratinocytes

Previous studies have used antibodies targeting VISTA to modulate cellular responses in the context of tumor models, sepsis, and psoriasis (*60–62*). To ask if modulating VISTA function *in vivo* suppresses UV-induced IFN-I response and skin injury, we treated humanized VISTA knock-in mice (hVISTA^KI^, expressing human VISTA in the mouse locus of the *Vsir* gene (*30*)) intraperitoneally with an anti-human VISTA antibody at the time of acute UV exposure (**Figure 6A**). hVISTA^KI^ mice that received anti-VISTA IgG showed reduced skin ISG expression 3 hours after UV and a ∼50% decrease in CLASI scores 48 hours after UV, compared to the isotype-treated controls (**Figure 6B-C).** Intradermal injection of anti-VISTA antibody 3 hours after UV also reduced ISG and IFN-κ expression one day after acute UV exposure (**Figure S6**). Therefore, antibody-mediated targeting of VISTA can effectively suppress UV-induced IFN-I and skin injury *in vivo*.

**Figure 6.**
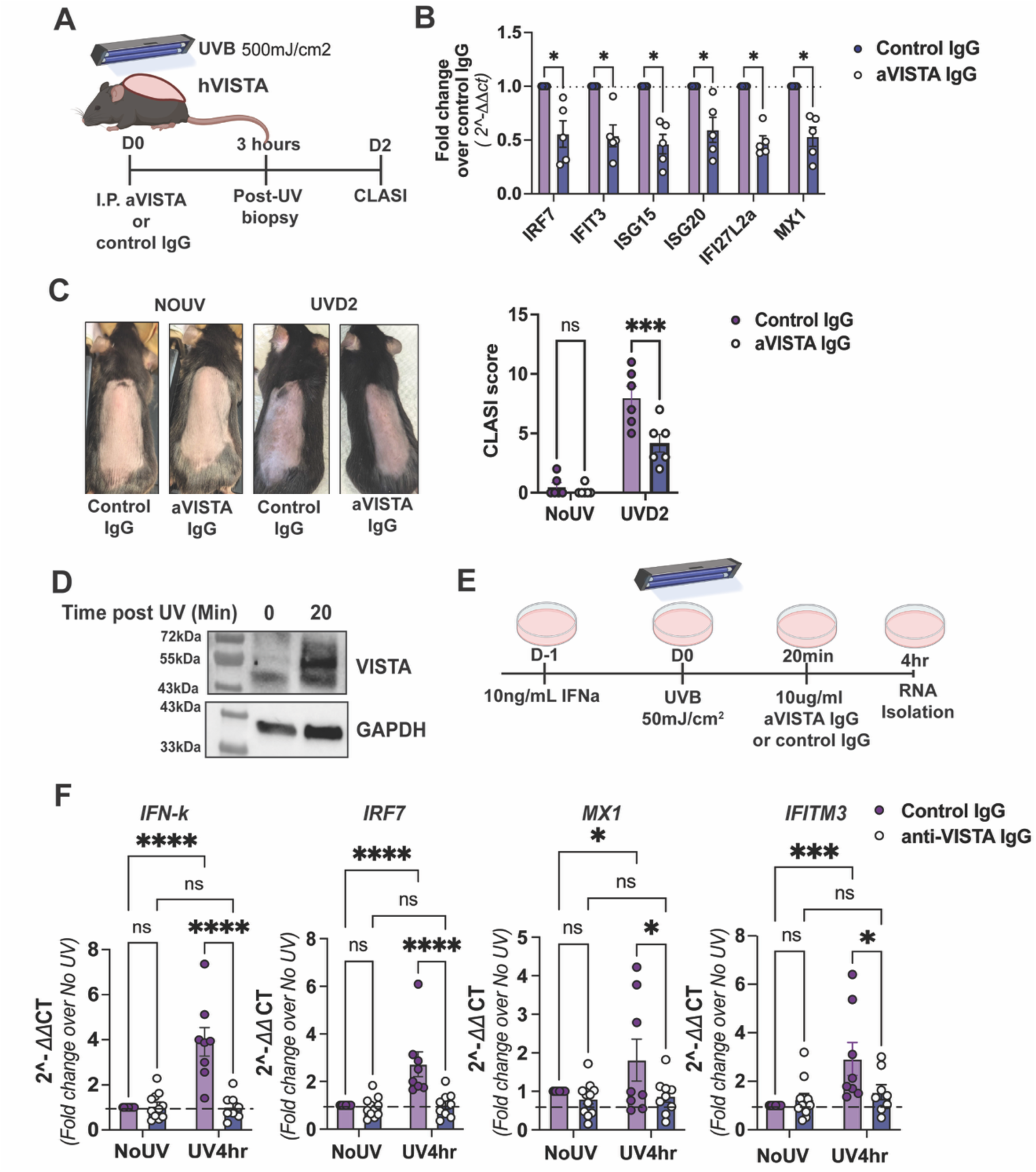
Anti-human VISTA antibody suppresses UV-induced injury and IFN-I response in the skin and epidermal keratinocytes. **(A)** Schematic of experimental design. Humanized VISTA-expressing mice (hVISTA^KI^) received anti-VISTA or isotype control IgG (i.p. 10ug/mL) immediately following exposure of the shaved dorsal back to UVB (500mJ/cm^2^). **(B)** Fold change in the expression of individual IFN-I stimulated genes (ISG), normalized to β*-actin*, in the skin of mice that received anti-VISTA or control IgG was compared with multiple Mann-Whitney tests (n=5, *p<0.05). **(C)** Representative images and CLASI scores of hVISTA^KI^ mice that received anti-VISTA or control IgG at the time of UV, before and 2 days post-UV were compared by two-way ANOVA with post-hoc Tukey test (n=6, ***p<0.001, ns = not significant). **(D)** Immunoblot for VISTA in human N/TERT keratinocytes before (0) and 20 minutes after UV *in vitro* (50mJ/cm^2^). **(E)** Schematic of experimental design. Human N/TERT keratinocytes were treated with IFNα (10ng/mL) for 16hr before exposure to UVB (50mJ/cm^2^) and treated with anti-VISTA IgG or isotype control (10ug/ml) 20min after UV. **(F)** Fold change in *IFN*κ and ISG (*IRF7, IFITM3, MX1*) transcript levels, normalized to *18S*, in N/TERT keratinocytes treated with anti-VISTA or control IgG after UV, relative to the levels in unexposed cells, were compared by two-way ANOVA with post-hoc Tukey test (n=8, * p<0.05, ***p<0.001,****p<0.0001, ns = not significant).

To translate the relevance of our findings in murine models to human disease, we investigated whether targeting VISTA could modulate UV-induced IFN-I and ISG expression in human keratinocytes pre-treated with IFN-α, simulating the IFN-I-rich environment of lupus skin. Interestingly, we detected high levels of VISTA in keratinocyte lysates by immunoblot just 20 minutes after exposure to UV (**Figure 6D**), which guided the timing of treatment with anti-VISTA IgG (**Figure 6E**). Keratinocytes treated with anti-VISTA IgG exhibited reduced expression of ISGs (*MX1, IFITM3, IRF7*) as well as *IFN-*κ 4 hours after UV (**Figure 6F**). Together, these findings suggest that targeting human VISTA may suppress UV-induced cutaneous IFN-I response both *in vivo* and in human keratinocytes under CLE-like conditions of high basal IFN-I.

## Discussion

The presented studies demonstrate that VISTA functions as a suppressor of cutaneous IFN-I response, both basally and following UVB exposure. Using young C57BL/6 mice, we demonstrate that *Vsir* deficiency leads to increased IFN-κ and ISG expression, resulting in a heightened skin IFN-I signature, as seen in non-lesional CLE skin. These data indicate that VISTA tonically limits cutaneous IFN-I independently of systemic autoimmunity, which is relevant to the observations that even non-lesional lupus skin shows an elevated baseline IFN-I signature (*4, 42*). Tonic IFN-I is necessary to maintain antiviral readiness (*25*), yet excessive IFN-I correlates with cutaneous disease activity in lupus (*63, 64*). Our results suggest that the loss of VISTA serves as a critical checkpoint in maintaining IFN-I homeostasis, and its loss lowers the threshold for inflammation, priming the skin for photosensitive pathology.

Keratinocyte-restricted VISTA loss (*K14^Cre^Vsir^fl/fl^*) mirrored the heightened skin IFN-I signature and UV-provoked IFN-I-mediated skin injury seen in global VISTA deficiency, identifying keratinocyte VISTA as essential for constraining skin IFN-I. These findings shine a light on a novel role for VISTA in keratinocytes, positioning epithelial cells as active regulators of cutaneous immune tone. These findings align with a growing body of literature emphasizing the importance of non-hematopoietic cells in lupus pathogenesis (*25, 65, 66*). VISTA is upregulated in bulge hair follicle stem cells (KC1) and basal (KC5) keratinocytes after UV. In the absence of VISTA, these keratinocyte subsets exhibited heightened IFN-I responses after UV exposure. The high expression of VISTA in follicular cells suggests a potential role for it in maintaining the hair follicle immune privilege and skin homeostasis. This is consistent with clinical observations that immune checkpoint blockade can trigger alopecia and rash (*67–70*) and prior work showing that VISTA homologue, PDL1, regulates hair cycling (*71*). Moreover, studies have demonstrated that follicular keratinocytes play a crucial role in the reepithelialization of the skin after wounding (*72*) and that UVB light damages the hair follicles of B6 mice (*73*). Indeed, we observed decreased expression of genes associated with hair cycle and ECM pathways in mouse keratinocytes deficient in VISTA, suggesting that keratinocyte VISTA may be involved in wound healing and repair after UV.

The effects of losing keratinocyte VISTA extended beyond the heightened skin IFN-I responses to a broader disruption of the basal skin environment. At homeostasis, K14^cre^*Vsir^fl^*^/fl^ skin has increased fibroblasts and T cells, but reduced keratinocytes and melanocytes, compared to cre-negative controls. Damage to melanocytes resulting in hypopigmentation is a common symptom of cutaneous lupus (*74, 75*). Importantly, keratinocytes provide many of the growth hormones that help maintain melanocytes, including Endothelin 2 and 3 (*Edn2, Edn3*), which were both decreased in K14^cre^*Vsir^fl^*^/fl^ keratinocytes at baseline. Since, Endothelin signaling is essential for melanocyte development (*76*), VISTA deficiency may disrupt an important keratinocyte-melanocyte axis. K14^cre^*Vsir^fl^*^/fl^ keratinocytes also express a variety of chemokines, stress markers, and MHC II molecules, which can be driven by exposure to type I and type II IFN (*77, 78*). Thus, loss of VISTA results in profoundly dysregulated keratinocytes that appear to be recruiting leukocytes and potentially presenting antigen.

The enhanced UV-induced injury in mice lacking keratinocyte VISTA is likely mediated by IFN-I driven recruitment and activation of inflammatory monocytes (*49*), which are elevated in the absence of keratinocyte VISTA and reduced when IFNAR is blocked. This supports recent studies identifying CD14+ monocytes as IFN-I-driven mediators of photosensitivity in human CLE (*79*). Monocyte-derived CD16+ dendritic cells have previously been identified as important intercellular communicators, primed to an inflammatory state by the IFN-I-rich environment of non-lesional lupus skin (*4, 80*). Future studies will investigate the fate of these inflammatory monocytes in mice to understand if they are differentiating into the pathogenic dendritic cells observed in lupus patients or whether they migrate to draining lymph nodes following UV, as recently reported (*81*).

In addition to its regulatory role in inflammation, our studies identify a role for keratinocyte VISTA in regulating skin barrier integrity and wound healing in response to UV. Keratinocytes from non-lesional lupus patients are inherently vulnerable to UV-induced apoptosis, likely due to chronic IFN-I exposure (*25*) or loss of epidermal growth factor receptor (EGFR) signaling (*82–84*), another hallmark of lupus skin. Recent work has implicated VISTA in suppressing EGFR trafficking/signaling in cancer cells (*85*) . UV exposure activates EGFR, which is expressed widely in keratinocytes, especially the hair follicle and suprabasal populations (*86, 87*). Thus, future investigations will test whether the VISTA-EGFR or VISTA-STING (elevated IFN-I) regulatory axes are crucial to keratinocyte homeostasis, differentiation and/or repair of UV-damaged skin.

While the studies presented here suggest VISTA may regulate IFN-I production by modulating STING levels and/or function, the regulatory mechanism at play warrants investigation in future studies. STING plays a central role in driving IFN-I production in response to endogenous nucleic acid accumulation (*20, 88*). STING gain-of-function mutations are a risk factor for SLE in humans (*89*), and recent reports support a critical role for STING in lupus pathogenesis, including photosensitivity (*13, 20*). Previous studies have proposed several molecular mechanisms by which VISTA may suppress intracellular inflammatory signaling, including regulation of TRAF6 polyubiquitination to dampen TLR signaling (*90*), promotion of innate lymphoid cell differentiation through FOXO1 transcriptional activity (*91*), and suppression of tumor cell growth via modulation of EGFR endosomal trafficking (*85*). STING itself is negatively regulated by recycling endosomal machinery (*92*), and IFN-I production and NF-κB responses downstream of STING are controlled by FOXO1 and TRAF6, respectively (*93, 94*). Therefore, future work to confirm the mechanistic link between VISTA and STING will test whether VISTA suppresses STING protein levels directly or IFN-I production and NF-κB signaling downstream of STING, or, alternatively, modulates cGAS activity via TRAF6 (*95*).

Previous studies have shown that antibody-mediated targeting of VISTA can suppress local and systemic features of disease in murine models of lupus and arthritis (*35, 96*). More recent studies have identified therapeutic benefits of targeting VISTA on specific cell population. For example, anti-VISTA IgG mitigated acute GVHD in models of hematopoietic stem cell transplantation (*29*) and limited K/BxN serum transfer arthritis (*96*) by suppressing T cells and neutrophils, respectively. In our studies, *in vivo* agonism of VISTA suppressed both UV-induced skin IFN-I response and subsequent skin injury, introducing a novel therapeutic application for VISTA targeting. Moreover, the findings presented here introduce a new VISTA-expressing cellular target, the epidermal keratinocytes, important contributors to CLE pathogenesis (*24, 25*). Keratinocytes express VISTA in response to UVB and VISTA activation on human keratinocytes *in vitro* suppresses UV-induced ISGs and IFN-κ. Therefore, targeting VISTA specifically in keratinocytes to modulate cell-intrinsic IFN-I production and/or response may be of therapeutic relevance to CLE. However, the mechanism by which anti-VISTA antibodies modulate keratinocyte function *in vivo* (e.g., by activating intrinsic VISTA or disrupting VISTA-ligand interactions) will be the subject of future work.

## Materials and Methods

### Mice and UV Irradiation

*Vsir-/- and Vsir^fl/fl^* mice were obtained from Dr. Randolph Noelle (refs). We crossed *Vsir^fl/fl^* mice with Tg(KRT14-cre)1Amc/J (K14-Cre, Strain: 004782, Jackson Laboratories) mice to generate mice with conditional deletion of keratinocyte VISTA (*K14^cre^Vsir^fl/f^)*. Controls for *Vsir-/-* mice were C57BL/6 mice from Charles River Laboratories (Strain Code: 027;Wilmington, MA). *Tmem173*-/- (STING KO mice, Strain: 025805) were purchased from Jackson Laboratories and crossed with *Vsir-/-* mice to obtain *Vsir-/- Tmem173-/-* mice. Human VISTA knock-in (hVISTA^KI^) were provided by Dr. Noelle (23).

Female 12–14-week-old mice were shaved dorsally with and electric razor and a thin layer of Nair^TM^ was applied by cotton swab for 30 seconds prior to removal by water. This procedure exposed about a 7.5cm^2^ area of back skin 5 days prior to UV exposure, to minimize the effects of shaving on skin inflammation. Mice were anesthetized with isoflurane and exposed to one dose of 500mJ/cm^2^ UVB using Tyler research UV-2 irradiation and dosimetry system. 250mJ/cm^2^ UVB dose was used for scRNA-seq experiments to ensure we captured more viable cells. Mice were given 3mg/kg ketoprofen and skin biopsies (6mm) were taken prior to and 3 hours after UVB. For measurement of UV-induced skin injury, mice were exposed to acute UVB and imaged before and 2 days post UVB. The back skin was scored for redness, scaling and scarring as described previously (*50*), in a blinded fashion, and scores summed into a cutaneous lupus and severity index (CLASI). All animals were maintained in a specific pathogen-free facility and studies were approved by Dartmouth’s Institutional Animal Care and Use Committee.

### Skin Bulk RNA-seq and qPCR

Total RNA from skin was isolated using the RNeasy Fibrous Tissue kit (Qiagen). cDNA was reverse transcribed using the iScript cDNA synthesis kit (Bio-rad) and quantitative PCR performed with iTaq Universal SYBR Green Supermix (Bio-rad). Primers for various ISGs and *Actin* are in Table 1. ISG expression CT values were normalized to *Actin*, and IFN-I scores were calculated as the sum of the normalized expression of 6 ISGs *(Irf7, Mx1, Ifi27l2a, Ifit1, Ifit3, Isg15* ) relative to baseline expression in the control skin as done in (*13*).

**Table 1:**
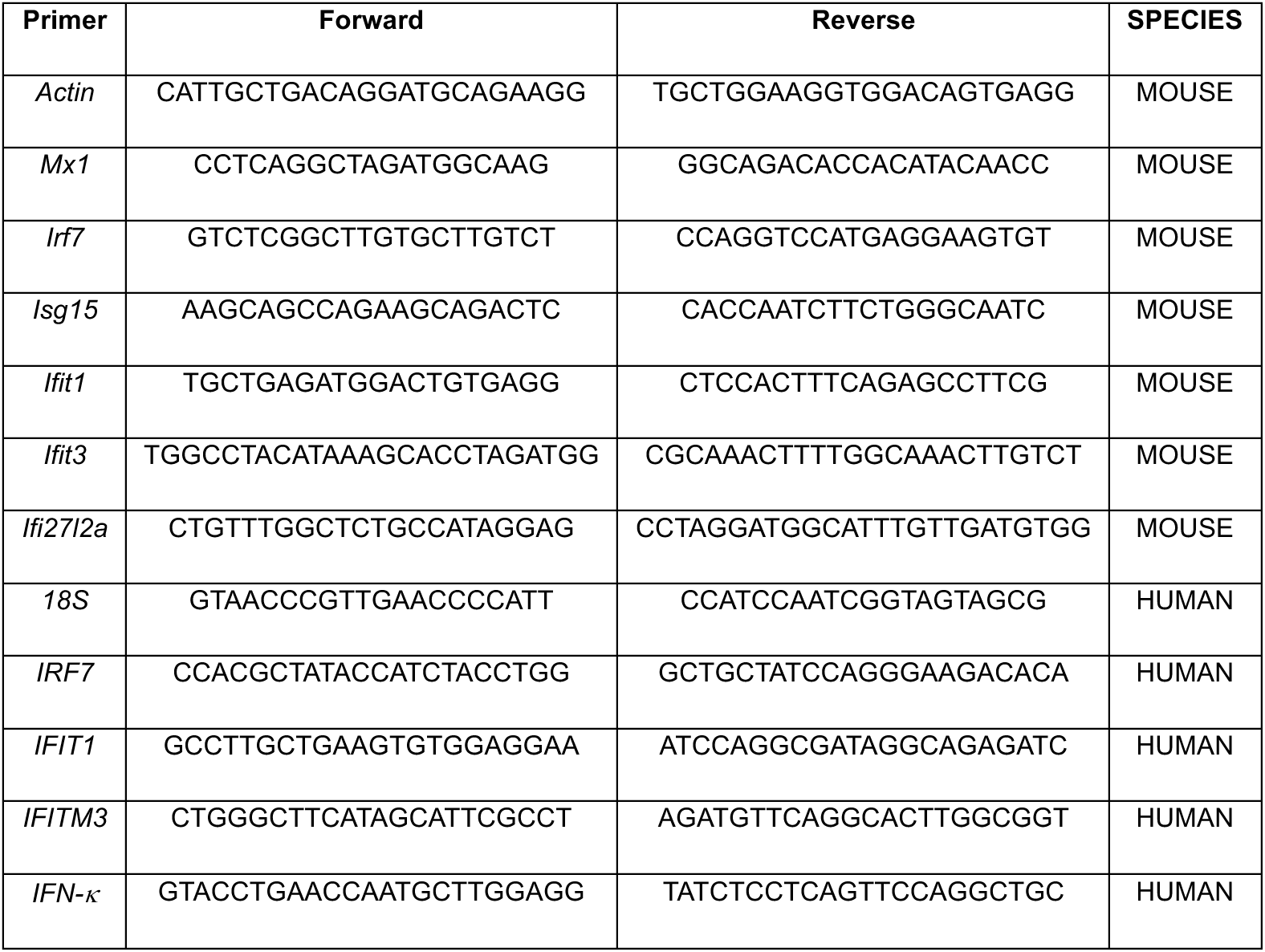
Primers Sequences for RT-qPCR.

For bulk RNA-seq, RNA quantity and integrity were assessed using a Qubit fluorometer (Thermo Fisher Scientific) and an Agilent Bioanalyzer 2100 (Agilent Technologies), respectively. Only samples with RNA integrity number (RIN) ≥ 8 were used for library preparation. rRNA depletion was performed using the Ribo-Zero Gold rRNA Removal Kit (Illumina) to enrich for coding and noncoding RNAs. cDNA libraries were prepared using the TruSeq Stranded Total RNA Library Prep Kit (Illumina) according to the manufacturer’s instructions. Sequencing was performed on an Illumina HiSeq 2000 platform to generate 100 bp paired-end reads, aiming for a depth of approximately [e.g., 30–50 million reads] per sample. RNA-sequencing data was analyzed using Deseq2 (V1.49) in R with pathway analysis performed in Webgestahlt.

### Human phototesting and Skin RNA sequencing

Healthy human and lupus patients from the University of Washington (UW) and Pennsylvania underwent phototesting (*13*). All individuals signed an informed consent in respective IRB-approved protocols (University of Washington; HSD number 50655). Briefly, participants were exposed to 2 minimally erythematous doses of UVB light and 6mm punch biopsies were obtained from unexposed skin and skin 24 hours after UV and stored in RNA later until RNA isolation at UW. cDNA was prepared from 1ug of total RNA and RNA-sequencing performed at the Northwest Genomics Center at the University of Washington as described previously (*13*).

### 10x Flex Genomics Single Cell RNA-sequencing

Formalin fixed paraffin embedded (FFPE) skin tissues blocks were prepared for the 10x Genomics Flex assay following demonstrated manufacturer’s protocol CG000606. For each block, 4 x 25um sections were placed into C tubes, deparaffinized with xylene, rehydrated with progressive dilutions of ethanol and dissociated in Liberase TM solution on a gentleMACS instrument (Miltenyi). The resulting cell/nuclei suspension was filtered through a 70um strainer and quantified using AO/PI staining on a Luna FX7 automated fluorescent cell counter.

The 10x Genomics Flex assay was performed according to the manufacturers protocol (CG000527). Briefly, cells were thawed from -80C, counted, and up to 1x106 cells were used as input into the hybridization reaction using the Mouse Transcriptome Probe Kit containing sample specific barcodes. Samples were pooled and loaded onto a 10x Genomics Chromium X instrument to capture 10,000 cells per sample. The concentration and size of sequencing libraries from the two pools were measured by Qubit (Thermo Fisher) and Tape Station (Agilent), respectively, and pooled for sequencing on a NextSeq2000 instrument (Illumina) to targeting 25,000 reads/cell. Raw fastq files were processed using the cellranger v9.0.0 pipeline prior to downstream analysis. Data were analyzed using Seurat (v5.3) Unsupervised clustering identified both immune and stromal clusters, which were identified by the top 5 expressed genes in each cluster. Sub- clustering of keratinocytes, macrophages, monocytes, and fibroblasts was performed and clusters identified by top 10 expressed genes. Differentially expressed genes and gene set enrichment analysis was performed using Deseq2 (1.48.2) and clusterProfiler (3.2.1). Ligand-receptor analysis of all identified clusters was performed by Cell Chat (V2.0).

### Mouse Keratinocyte Isolation

Mice were euthanized with CO2, tails removed at the base, and keratinocytes isolated as published (*97*). Briefly, tails were rinsed with PBS in a petri dish and a scalpel used to cut along the tail skin from base to tip. Skin was peeled away and suspended into Epilife with 4ug/mL dispase (Gibco) and rotated for 14-16 hours at 4°C. Tail skin was washed in PBS and epidermis peeled off with forceps. Epidermis was placed basal side down on 4 mL 0.05% trypsin and incubated with gentle shaking at 37°C for 20 minutes. 10 mL Epilife with 10% FBS was added to quench trypsin reaction, epidermis was scrubbed against petri dish to free keratinocytes, and cells were filtered through 100- micron filters twice for a total of 30 mLs Cells were centrifuged at 180xG for 5 minutes, resuspended in 5 mLs Epilife with penicillin/streptomycin, anti-mycotic/antibiotic and human keratinocyte growth supplement, and seeded at 1x10^5^ x cm^2^ density. After 24 hours, media was removed and cells washed gently with PBS before relacing with fresh media. Cells were then grown with media changed every 2-3 days until confluency.

### Histology

Tissue sections from back skin of mice were fixed in 10% Neutral Buffered Formalin for 24-48 hrs then embedded in paraffin. 4um sections were stained following the guidelines in the Opal 7-color Manual IHC Kit (Akoya Biosciences, Catalog: NEL811001KT). Antibodies used for staining include anti-VISTA (D5L5T, CST: 54979) and anti-CK14 (Invitrogen, MA5-11599). Nuclei were stained with 1X Dapi [Invitrogen, D1306] and mounted with Prolong Diamond (Thermo Fisher, P36961). A Vectra 3.0 microscope was used for image acquisition. Briefly, whole slide scans were done, Phenochart was used to select MSI regions, and images were acquired at 40x magnification. Spectral libraries were generated using an unstained control slide, single color control slides for each Opal stain, and a slide for DAPI. Click-iT™ Plus TUNEL Assay [Invitrogen, C10619] was performed according to the manufacturer’s protocol. Tissues were visualized using the Keyence BZ-X800 fluorescent microscope.

### Immunoblots

Cells or skin biopsies were snap frozen at -80C or in liquid nitrogen respectively and lysed in RIPA buffer after thawing (or PBS for homogenates) with Halt protease inhibitor. Biopsies were additionally subjected to bead sonication. Protein concentrations were determined by Bradford assay. Lysates were diluted in Lammeli buffer with DTT (50mM), boiled for 5 minutes at 95°C and then centrifuged at 10,000 RPM for 5 minutes. 15-40ug of protein were resolved on 12% polyacrylamide SDS-PAGE at 150V for 1 hour and transferred to a nitrocellulose membrane at 100V for 1 hour. Membranes were blocked in 5% milk for 1 hr and incubated overnight in primary antibody: anti-mouse IFN-κ (AF5206), anti-mouse β-ACTIN (CST, 4967S), anti-human/mouse GAPDH (Cell Signaling, 3700S). Antibody dilutions are described in Supplemental Table 2.

### Enzyme-Linked Immunosorbent Assay (ELISA)

Quantification of IFN-κ levels in mouse skin homogenates were performed using a commercially available ELISA kit (DFNAS0; Mybiosource:, MBS287021) according to the manufacturer’s instructions. Briefly, 40ug of protein from skin homogenates was loaded evenly into wells containing capture antibodies and incubated for 1-2 hours. Plates were then washed 3 times following incubation with detection antibody, washed again and then incubated with streptavidin-HRP. Plates were washed a final time before incubation with substrate and stopping solution. Plates were read on a BioTek Synergy HTX at 405nm and 450nm (BioTek, 0211-3030).

### Anti-human VISTA antibody treatments *in vitro* and *in vivo*

Human N/TERT keratinocytes were cultured in keratinocyte SFM media containing 30ug/mL BPE and 0.2ng/mL EGF, 310uM CaCL^2+^. Media was changed ever 2 days and cells grown until 50% confluent for passaging and 70% confluency for experiments. Cell were treated with anti-VISTA IgG (INX803, 10ug/mL) 20 minutes after UVB. RNA was isolated using Zymogen total RNA isolation kit 4hr after UVB. For *in vivo* treatment of humanized VISTA mice, animals were anesthetized, exposed to UVB (500mJ/cm^2^) and immediately injected intra peritoneally or intradermally with 10ug of anti-VISTA GG8 or INX803 antibody or isotype control. IFN-I and ISG transcript levels were quantified 3 hours post UVB in whole skin biopsies by qPCR. CLASI scores and immune infiltration were quantified in unirradiated and mice 2 days after UVB.

### Flow Cytometry

Mouse ears were excised, split and incubated dermis side down in Liberase TL (0.25mg/mL) and DNAse I (0.5mg/mL) for 1.5 hours at 37°C with shaking. Ears were then chopped with microscissors and added to gentleMACS C tubes (Miltenyi Biotech) and mechanically dissociated with program Multi_H_C_Tube. After passing through a 70uM strainer and washing, cells were stained with Live Dead blue viability dye (ThermoFisher Scientific). Samples were incubated with TruStain FcX PLUS (S170011E; Biolegend) and stained with antibody cocktail. Flow cytometry was performed on a Cytek Aurora instrument. Data was analyzed using FlowJo software (Treestar Incorporated, Ashland, OR) following initial gating on Live/Dead negative singlets. The following antibodies: CD45, CD11b, Ly6C, Ly6G, F480 were purchased conjugated to APC-Cy7, Percep_Cy5.5, BV510, Spark YG 593, BV421 from Biolegend (San Diego, CA), ThermoFisher Scientific (Waltham, MA), or BD Biosciences (Franklin Lakes, NJ).

### Reanalysis of published single-cell RNA-seq dataset

Publicly available human single-cell RNA-sequencing data from 14 healthy control and 7 non-lesional lupus patient skin samples were reanalyzed using Seurat (v5.3) identifying keratinocyte populations and annotated using the top 5 expressed genes in each subcluster. Expression of *VSIR* and GO:Response to IFN-I score was quantified across clusters by condition.

## Acknowledgments

This work was also supported by Lupus Research Alliance, CDMRP Lupus Program, NIAMS (R21 AR079661-01), and Dartmouth Cancer Center Prouty award to S.S.-G; NIAID R01 AI148430-05 to S.S.G. and R.M.; NIAMS P30-AR075043 for J.E.G.; NHLBI R01 HL155144-04 for B.R.B. This work was performed with the assistance of the Immune Monitoring and Flow Cytometry Resource, Irradiation, Pre-clinical Imaging and Microscopy Resource, and the Pathology Shared Resource. RNA-sequencing was carried out in the Genomics and Molecular Biology Shared Resource (RRID:SCR_021293) at Dartmouth which is supported by NCI Cancer Center Support Grant 5P30CA023108 and NIH S10 (1S10OD030242) awards and single-cell studies were conducted through the Dartmouth Center for Quantitative Biology in collaboration with the GMBSR with support from NIGMS (P20GM130454) and NIH S10 (S10OD025235) awards.

---

**Figure S1:**
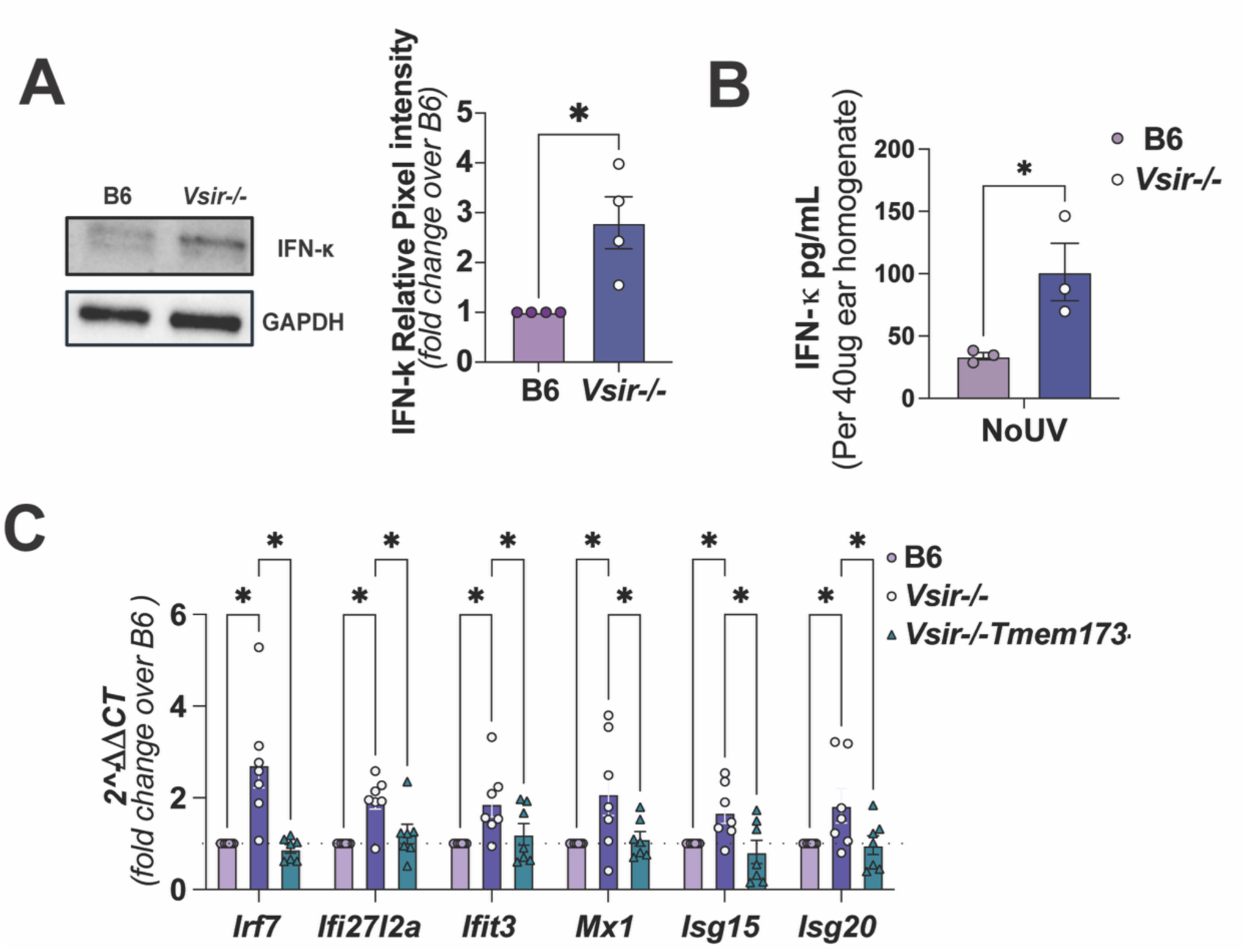
IFN. κ **and ISG levels in the skin of VISTA-deficient mice. (A)** Representative IFNk immunoblot of B6 and *Vsir-/-* mouse skin and ImageJ quantification relative to GAPDH compared by Mann-Whitney test (n=5; *p<0.05). **(B)** IFN-k levels in B6 and *Vsir-*/- mouse skin homogenates quantified by ELISA; significance was determined by independent t-test (n=5; *p<0.05). **(C)** Fold change in basal ISGs in B6, *Vsir-*/- and *Vsir-*/-*Tmem173*-/- mouse skin was compared using One-way ANOVA with post-hoc Tukey test (*Irf7, Ifi27l2a, Ifit3, Mx1, Isg15, Isg20)* to *18s*, in *Vsir*-/- and *Vsir*-/-*Tmem173-/-* relative to B6 skin.

**Figure S2.**
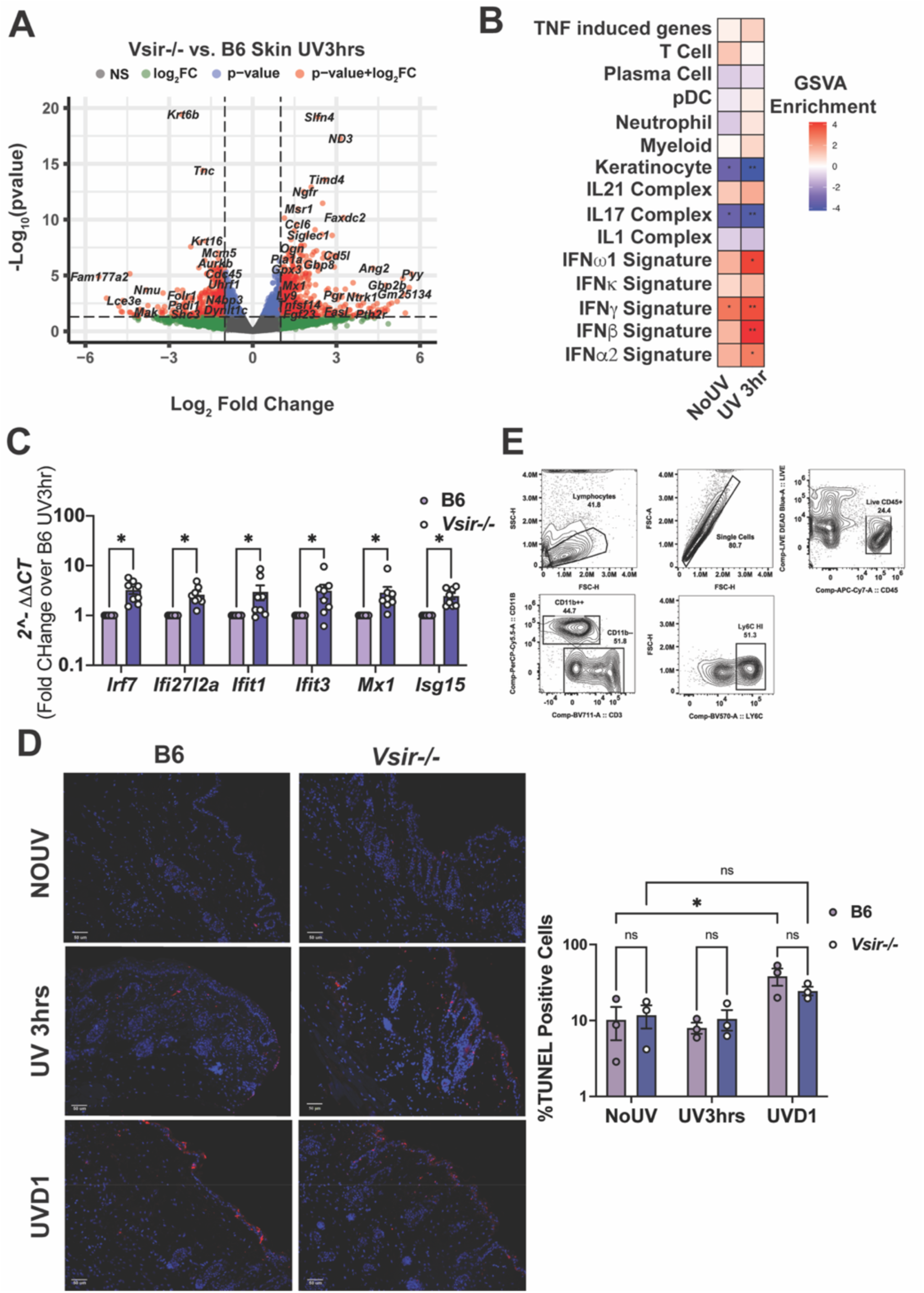
UV-induced IFN-I response is elevated in *Vsir-/-* mouse skin. **(A)** Differentially expressed genes in VISTA-deficient (*Vsir*-/-) mouse compared to wild-type B6 skin, 3 hours post UVB (FDR adjusted p value < 0.01 and -1 < Log2FC > 1). **(B)** Comparative gene set variation analysis (GSVA) showing hedges G effect size of pathway enrichment in *Vsir-/-* skin before and 3 hours post-UVB compared to B6 skin by Welsh’s t-test (n=3; *p<0.05, **p<0.01). **(C)** Fold change in ISG expression in *Vsir-/-* vs. B6 mouse skin 3 hours post UVB exposure (500mJ/cm^2^) compared by multiple Mann-Whitney tests, controlled for false discovery by Benjamini, Krieger Yekutieli test (n=8, *p<0.05). **(D)** TUNEL staining of B6 and *Vsir-/-* skin at baseline, 3 hours, and 1 day post-UV and quantification of TUNEL+ cells by ImageJ determined by two-way ANOVA with post-hoc Tukey test (n=3; *p<0.05, ns = not significant)**. (E)** Representative gating strategy for identifying Ly6C^hi^ monocytes in mouse skin.

**Figure S3.**
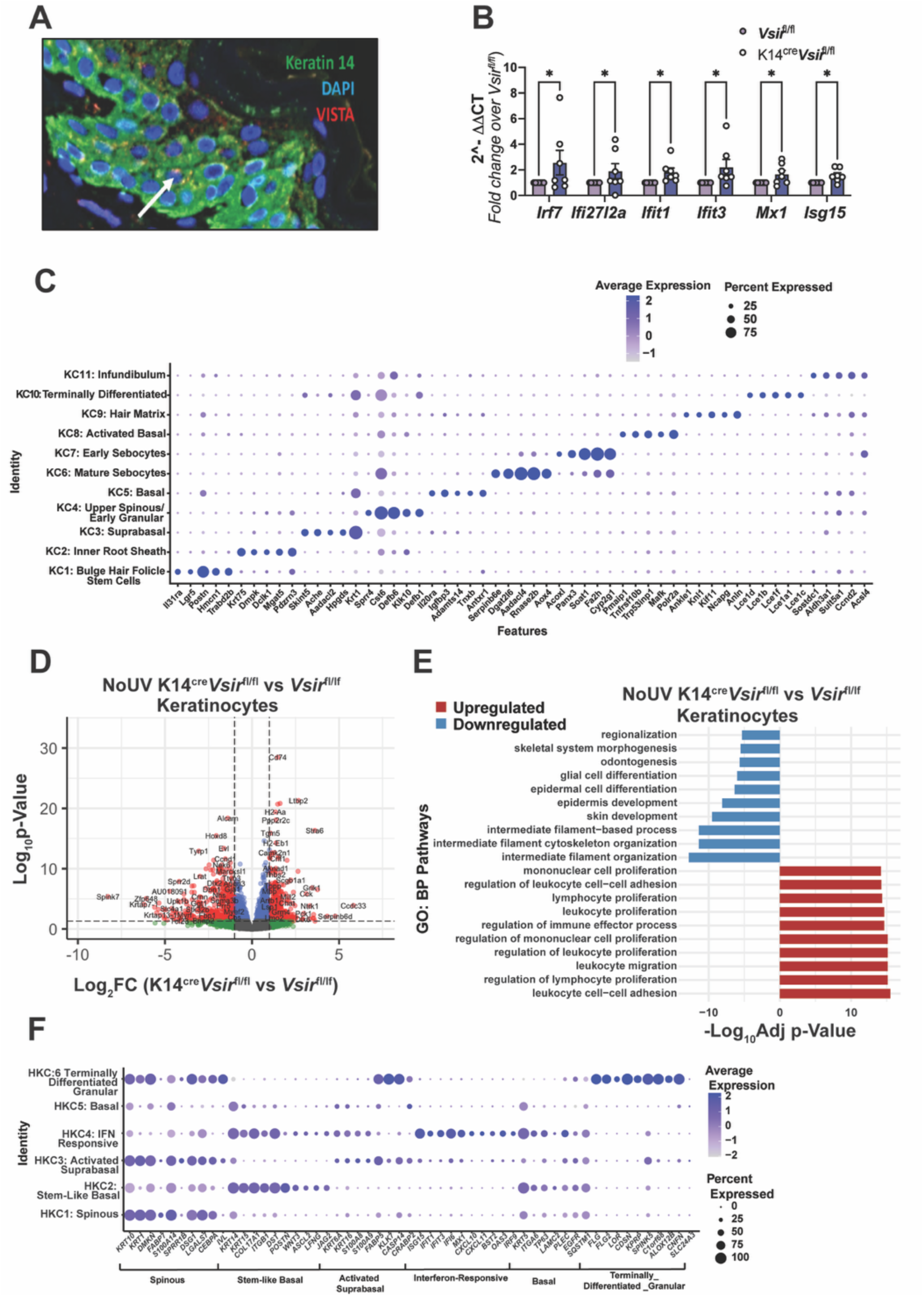
Loss of keratinocyte VISTA disrupts keratinocyte differentiation and skews keratinocyte subsets toward an activated and antigen-presenting state. **(A)** Mouse epidermis stained for VISTA (red), Keratin-14 (green), and DAPI (blue). **(B)** Fold change in ISGs in K14^Cre^*Vsir^fl^*^/fl^ skin at baseline compared to *Vsir^fl^*^/fl^ skin by multiple Mann- Whitney tests, corrected with Benjamini, Krieger Yekutieli test (n=7, *p<0.05). **(C)** Dot plot showing top 10 signature genes in each keratinocyte cluster from K14^Cre^*Vsir^fl^*^/fl^ and *Vsir^fl^*^/fl^ mouse keratinocytes from both conditions (No UV and 24hr after UV). Dot size represents the percent of subcluster expressed in each cell and color demonstrates normalized expression of each gene. **(D)** Differentially expressed genes in K14^Cre^*Vsir^fl^*^/fl^ vs *Vsir^fl^*^/fl^ keratinocyte cluster at baseline (n = 4 per genotype). **(E)** Top 10 upregulated and downregulated pathways (GO: Biological Processes) from select IFN-I rich K14^Cre^*Vsir*^fl/fl^ vs *Vsir*^fl/fl^ baseline keratinocyte subclusters. Enrichment scores are normalized to the size of the gene set, and p-values are determined by permutation testing and multiple comparisons corrected for by Benjamini-Hochberg (n=4). **(F)** Top 10 expressed genes across human keratinocyte subsets were selected based on gene set enrichment analysis (GSE186476).

**Figure S4.**
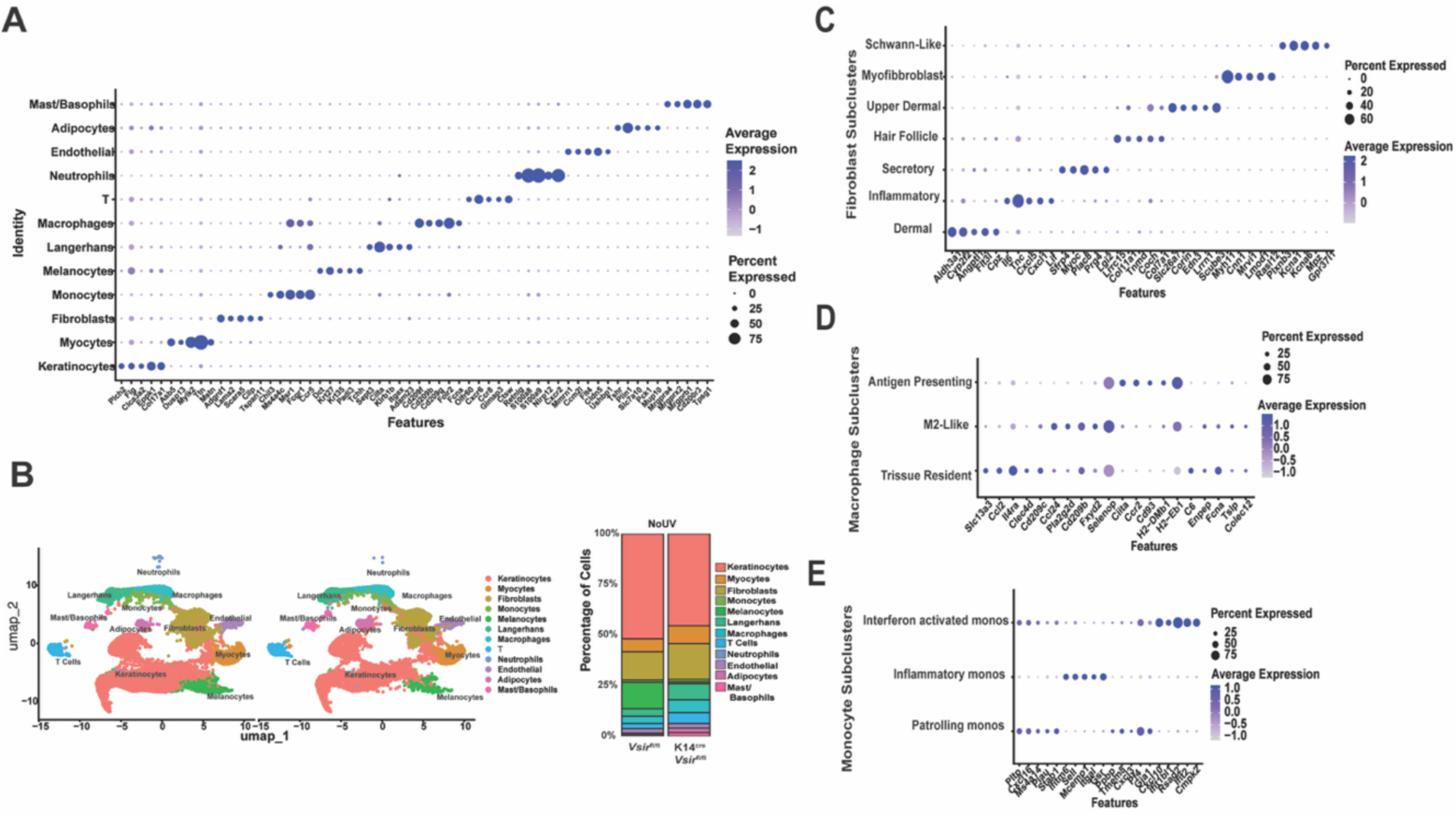
Gene signatures define skin immune and stromal cells in the absence of keratinocyte VISTA. **(A)** Top 5 expressed genes across immune cell clusters were selected based on gene set enrichment analysis. (**B)** UMAP of baseline K14^Cre^*Vsir*^fl/fl^ and *Vsir*^fl/fl^ skin cell clusters, split by genotype; relative percentage of each cluster is represented in bar graphs. **(C-E)** Top 5 expressed genes across clusters within (**C**) fibroblasts, (**D**) macrophages, and (**E**) monocytes were selected based on gene set enrichment analysis. Dot size represents the percent of subcluster expressed in each cell and color demonstrates normalized expression of each gene.

**Figure S5.**
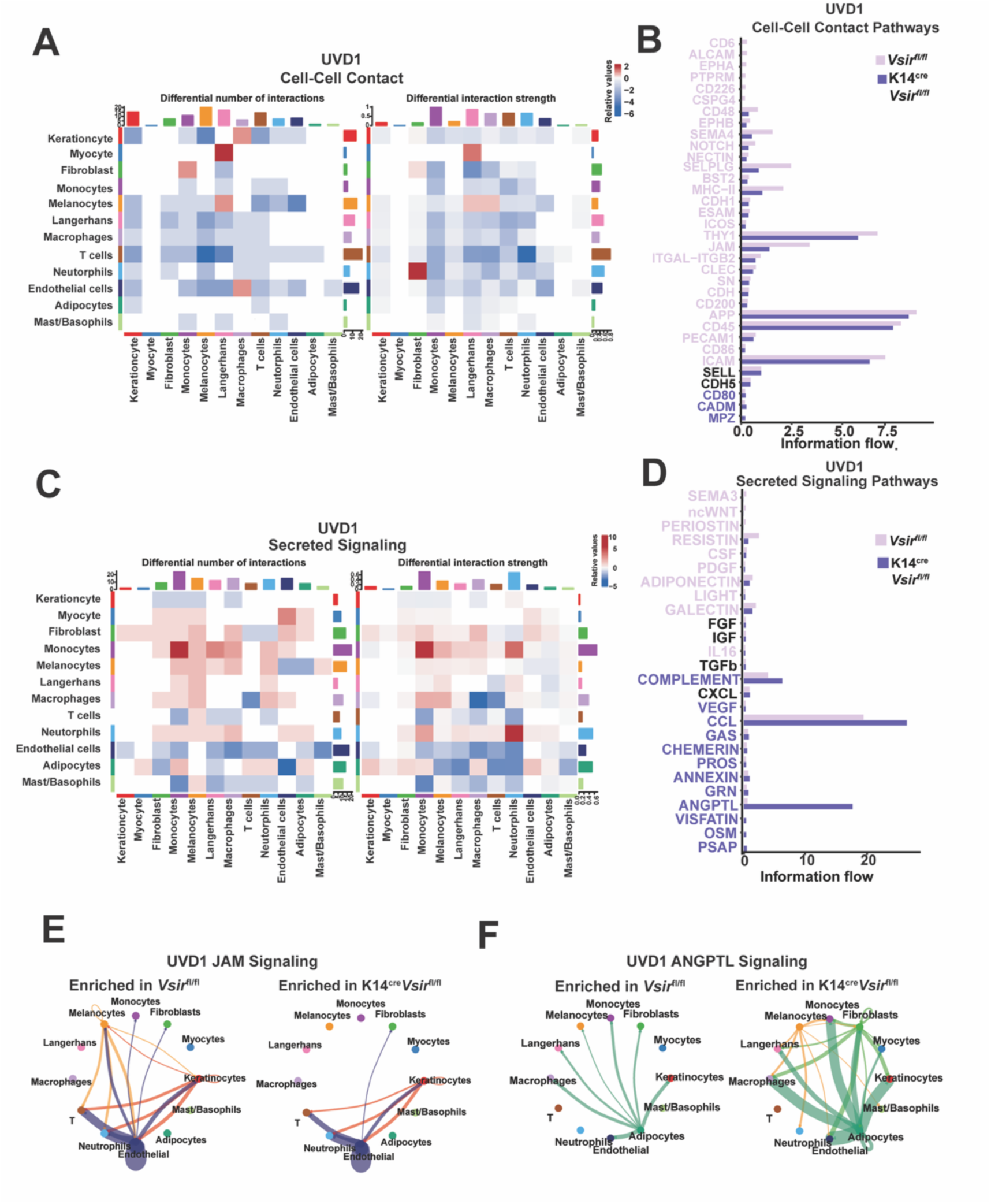
Keratinocyte VISTA deficiency alters cell-cell communication in the skin after UV exposure. Ligand receptor analysis (Cell-Chat) showing changes in the number and strength of **(A-B)** Cell-Cell contact and **(C-D)** secreted signaling interactions in K14^Cre^*Vsir^fl^*^/fl^ compared to *Vsir^fl^*^/fl^ skin, 1 day after UV. Interaction strength is calculated as average expression of the sender (ligand) multiplied by receiver (receptor) genes. (**B,D**) Information flow of **(B)** cell-cell contact and **(D)** secreted signaling pathways in K14^Cre^*Vsir^fl^*^/fl^ skin 1 day after UV. Overall information flow is calculated as the sum of probabilities (edges) that denote an interaction between cell types and significant differences in pathway information flow are determined by permutation testing. Pathways upregulated in K14^Cre^*Vsir^fl^*^/fl^ are labeled in purple and pathways decreased in K14^Cre^*Vsir^fl^*^/fl^ are labeled in pink. **(E-F)** Circos plots showing (E) change in cell-cell contact signaling pathway JAM and (F) secreted signaling pathway ANGPTL in K14^Cre^*Vsir^fl^*^/fl^ vs *Vsir^fl^*^/fl^ skin cells 1 day after UV.

**Figure S6.**
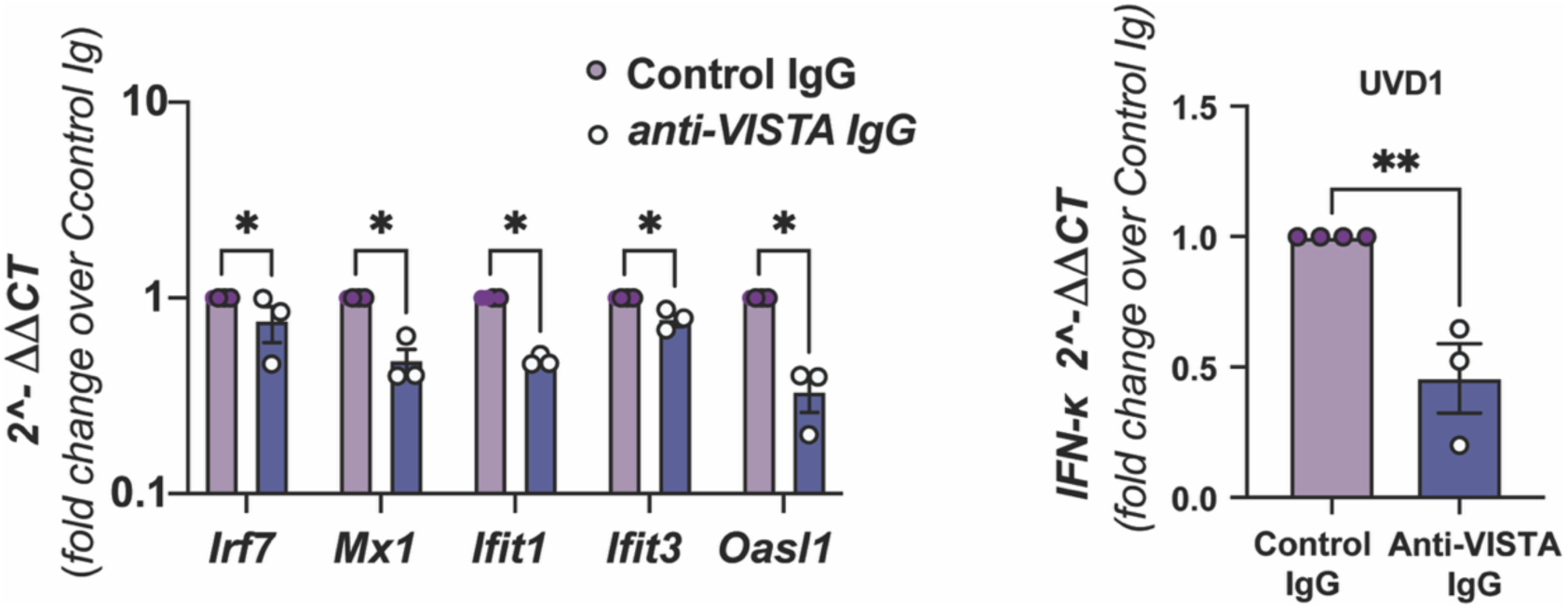
Intradermal injection of anti-VISTA antibody suppresses UV-induced IFN- I response in the skin. Fold change in the expression of ISGs and *Ifn-*κ normalized to *18s*, in the skin of mice that received anti-VISTA or control IgG was compared with multiple Mann-Whitney tests (n=3 Control IgG; n=4 anti-VISTA IgG, *p<0.05.

## Notes

### Competing Interest Statement

The authors have declared no competing interest.

## References

1. D. Wong et al., Interferon and biologic signatures in dermatomyositis skin: specificity and heterogeneity across diseases. PLoS One 7, e29161 (2012).

2. X. Wang et al., Identification and validation of interferon-stimulated gene 15 as a biomarker for dermatomyositis by integrated bioinformatics analysis and machine learning. Front Immunol 15, 1429817 (2024).

3. T. Miyamoto et al., Assessment of type I interferon signatures in undifferentiated inflammatory diseases: A Japanese multicenter experience. Front Immunol 13, 905960 (2022).

4. A. C. Billi et al., Nonlesional lupus skin contributes to inflammatory education of myeloid cells and primes for cutaneous inflammation. Sci Transl Med 14, eabn2263 (2022).

5. I. Braunstein, R. Klein, J. Okawa, V. P. Werth, The interferon-regulated gene signature is elevated in subacute cutaneous lupus erythematosus and discoid lupus erythematosus and correlates with the cutaneous lupus area and severity index score. Br J Dermatol 166, 971–975 (2012).

6. V. Pascual, L. Farkas, J. Banchereau, Systemic lupus erythematosus: all roads lead to type I interferons. Curr Opin Immunol 18, 676–682 (2006).

7. R. H. Refai, M. F. Hussein, M. H. Abdou, A. N. Abou-Raya, Environmental risk factors of systemic lupus erythematosus: a case-control study. Sci Rep 13, 10219 (2023).

8. B. Klein, M. Kunz, Current concepts of photosensitivity in cutaneous lupus erythematosus. Front Med (Lausanne) 9, 939594 (2022).

9. H. Wolska, M. Blaszczyk, S. Jablonska, Phototests in patients with various forms of lupus erythematosus. Int J Dermatol 28, 98–103 (1989).

10. A. Kuhn et al., Phototesting in lupus erythematosus: a 15-year experience. J Am Acad Dermatol 45, 86–95 (2001).

11. F. Furukawa et al., Keratinocytes from patients with lupus erythematosus show enhanced cytotoxicity to ultraviolet radiation and to antibody-mediated cytotoxicity. Clin Exp Immunol 118, 164–170 (1999).

12. A. Kim, B. F. Chong, Photosensitivity in cutaneous lupus erythematosus. Photodermatol Photoimmunol Photomed 29, 4–11 (2013).

13. S. Skopelja-Gardner et al., The early local and systemic Type I interferon responses to ultraviolet B light exposure are cGAS dependent. Sci Rep 10, 7908 (2020).

14. J. He, Y. Guo, J. Chen, J. Xu, X. Zhu, Exploring the correlation between UVB sensitivity and SLE activity: Insights into UVB-driven pathogenesis in lupus erythematosus. J Autoimmun 153, 103393 (2025).

15. E. F. Morand et al., Trial of Anifrolumab in Active Systemic Lupus Erythematosus. New England Journal of Medicine 382, 211–221 (2020).

16. E. D. Papadimitraki, G. K. Bertsias, D. T. Boumpas, Toll like receptors and autoimmunity: a critical appraisal. J Autoimmun 29, 310–318 (2007).

17. C. G. Horton, Z. J. Pan, A. D. Farris, Targeting Toll-like receptors for treatment of SLE. Mediators Inflamm 2010, (2010).

18. L. E. Munoz, K. Lauber, M. Schiller, A. A. Manfredi, M. Herrmann, The role of defective clearance of apoptotic cells in systemic autoimmunity. Nat Rev Rheumatol 6, 280–289 (2010).

19. M. Shrivastav, T. B. Niewold, Nucleic Acid sensors and type I interferon production in systemic lupus erythematosus. Front Immunol 4, 319 (2013).

20. P. P. Vejvisithsakul et al., Elucidating the function of STING in systemic lupus erythematosus through the STING Goldenticket mouse mutant. Sci Rep 14, 13968 (2024).

21. L. C. Tsoi et al., Hypersensitive IFN Responses in Lupus Keratinocytes Reveal Key Mechanistic Determinants in Cutaneous Lupus. J Immunol 202, 2121–2130 (2019).

22. S. Song et al., Inhibition of IRF5 hyperactivation protects from lupus onset and severity. J Clin Invest 130, 6700–6717 (2020).

23. D. Li, et al., IRF5 genetic risk variants drive myeloid-specific IRF5 hyperactivation and presymptomatic1 SLE. JCI Insight 5, (2020).

24. B. Klein et al., Epidermal ZBP1 stabilizes mitochondrial Z-DNA to drive UV-induced IFN signaling in autoimmune photosensitivity. Sci Immunol 10, eado1710 (2025).

25. M. K. Sarkar et al., Photosensitivity and type I IFN responses in cutaneous lupus are driven by epidermal-derived interferon kappa. Annals of the Rheumatic Diseases 77, 1653–1664 (2018).

26. B. Klein et al., Epidermal Interferon kappa drives cutaneous lupus-like lesions, photosensitivity and systemic autoimmunity in vivo. Arthritis Rheumatol, (2025).

27. L. Wang et al., VISTA, a novel mouse Ig superfamily ligand that negatively regulates T cell responses. J Exp Med 208, 577–592 (2011).

28. M. A. ElTanbouly, W. Croteau, R. J. Noelle, J. L. Lines, VISTA: a novel immunotherapy target for normalizing innate and adaptive immunity. Semin Immunol 42, 101308 (2019).

29. B. H. Koehn et al., Targeting cell-surface VISTA expression on allospecific naive T cells promotes tolerance. Blood 145, 1687–1700 (2025).

30. M. A. ElTanbouly et al., VISTA is a checkpoint regulator for naive T cell quiescence and peripheral tolerance. Science 367, (2020).

31. L. Wang et al., Programmed death 1 ligand signaling regulates the generation of adaptive Foxp3+CD4+ regulatory T cells. Proc Natl Acad Sci U S A 105, 9331–9336 (2008).

32. M. A. ElTanbouly et al., VISTA Re-programs Macrophage Biology Through the Combined Regulation of Tolerance and Anti-inflammatory Pathways. Front Immunol 11, 580187 (2020).

33. Y. Lin et al., VISTA drives macrophages towards a pro-tumoral phenotype that promotes cancer cell phagocytosis yet down-regulates T cell responses. Exp Hematol Oncol 13, 35 (2024).

34. M. A. ElTanbouly, E. Schaafsma, R. J. Noelle, J. L. Lines, VISTA: Coming of age as a multi- lineage immune checkpoint. Clin Exp Immunol 200, 120–130 (2020).

35. X. Han et al., PD-1H (VISTA)-mediated suppression of autoimmunity in systemic and cutaneous lupus erythematosus. Sci Transl Med 11, (2019).

36. B. Ersek et al., Melanoma-associated fibroblasts impair CD8+ T cell function and modify expression of immune checkpoint regulators via increased arginase activity. Cell Mol Life Sci 78, 661–673 (2021).

37. S. Ceeraz et al., VISTA Deficiency Accelerates the Development of Fatal Murine Lupus Nephritis. Arthritis Rheumatol 69, 814–825 (2017).

38. K. W. Yoon et al., Control of signaling-mediated clearance of apoptotic cells by the tumor suppressor p53. Science 349, 1261669 (2015).

39. P. A. Sergent et al., Blocking the VISTA pathway enhances disease progression in (NZB × NZW) F1 female mice. Lupus 27, 210–216 (2018).

40. S. Schlichtner et al., Expression of the Immune Checkpoint Protein VISTA Is Differentially Regulated by the TGF-beta1 - Smad3 Signaling Pathway in Rapidly Proliferating Human Cells and T Lymphocytes. Front Med (Lausanne) 9, 790995 (2022).

41. S. R. Rosenbaum et al., FOXD3 Regulates VISTA Expression in Melanoma. Cell Reports 30, 510–524.e516 (2020).

42. T. M. Li et al., The interferon-rich skin environment regulates Langerhans cell ADAM17 to promote photosensitivity in lupus. Elife 13, (2024).

43. B. A. Martinez et al., Machine learning reveals distinct gene signature profiles in lesional and nonlesional regions of inflammatory skin diseases. Sci Adv 8, eabn4776 (2022).

44. J. Patel et al., Multidimensional Immune Profiling of Cutaneous Lupus Erythematosus In Vivo Stratified by Patient Response to Antimalarials. Arthritis Rheumatol 74, 1687–1698 (2022).

45. J. N. Stannard et al., Lupus Skin Is Primed for IL-6 Inflammatory Responses through a Keratinocyte-Mediated Autocrine Type I Interferon Loop. J Invest Dermatol 137, 115– 122 (2017).

46. O. Demaria, J. Di Domizio, M. Gilliet, Immune sensing of nucleic acids in inflammatory skin diseases. Semin Immunopathol 36, 519–529 (2014).

47. H. Wobma, D. S. Shin, J. Chou, F. Dedeoglu, Dysregulation of the cGAS-STING Pathway in Monogenic Autoinflammation and Lupus. Front Immunol 13, 905109 (2022).

48. S. J. Wolf et al., Ultraviolet light induces increased T cell activation in lupus-prone mice via type I IFN-dependent inhibition of T regulatory cells. J Autoimmun 103, 102291 (2019).

49. C. Sontheimer, D. Liggitt, K. B. Elkon, Ultraviolet B Irradiation Causes Stimulator of Interferon Genes-Dependent Production of Protective Type I Interferon in Mouse Skin by Recruited Inflammatory Monocytes. Arthritis Rheumatol 69, 826–836 (2017).

50. J. Albrecht et al., The CLASI (Cutaneous Lupus Erythematosus Disease Area and Severity Index): an outcome instrument for cutaneous lupus erythematosus. J Invest Dermatol 125, 889–894 (2005).

51. X. Huang et al., VISTA: an immune regulatory protein checking tumor and immune cells in cancer immunotherapy. J Hematol Oncol 13, 83 (2020).

52. M. Borggrewe et al., VISTA expression by microglia decreases during inflammation and is differentially regulated in CNS diseases. Glia 66, 2645–2658 (2018).

53. S. J. Luk et al., VISTA Expression on Cancer-Associated Endothelium Selectively Prevents T-cell Extravasation. Cancer Immunol Res 11, 1480–1492 (2023).

54. H. R. Dassule, P. Lewis, M. Bei, R. Maas, A. P. McMahon, Sonic hedgehog regulates growth and morphogenesis of the tooth. Development 127, 4775–4785 (2000).

55. C. Li et al., DNA damage-triggered activation of cGAS-STING pathway induces apoptosis in human keratinocyte HaCaT cells. Mol Immunol 131, 180–190 (2021).

56. A. J. Sanders, D. G. Jiang, W. G. Jiang, K. G. Harding, G. K. Patel, Activated leukocyte cell adhesion molecule impacts on clinical wound healing and inhibits HaCaT migration. Int Wound J 8, 500–507 (2011).

57. X. Huang et al., CDH1 is Identified as A Therapeutic Target for Skin Regeneration after Mechanical Loading. Int J Biol Sci 17, 353–367 (2021).

58. C. Siemes et al., Keratinocytes from APP/APLP2-deficient mice are impaired in proliferation, adhesion and migration in vitro. Exp Cell Res 312, 1939–1949 (2006).

59. Y. Wang et al., JAM-A knockdown accelerates the proliferation and migration of human keratinocytes, and improves wound healing in rats via FAK/Erk signaling. Cell Death Dis 9, 848 (2018).

60. Y. V. Ma, A. Sparkes, E. Romao, S. Saha, J. Gariepy, Agonistic nanobodies and antibodies to human VISTA. MAbs 13, 2003281 (2021).

61. S. Iadonato et al., A highly potent anti-VISTA antibody KVA12123 - a new immune checkpoint inhibitor and a promising therapy against poorly immunogenic tumors. Front Immunol 14, 1311658 (2023).

62. T. Tao et al., High-Affinity Anti-VISTA Antibody Protects against Sepsis by Inhibition of T Lymphocyte Apoptosis and Suppression of the Inflammatory Response. Mediators Inflamm 2021, 6650329 (2021).

63. L. Mai et al., The baseline interferon signature predicts disease severity over the subsequent 5 years in systemic lupus erythematosus. Arthritis Res Ther 23, 29 (2021).

64. E. Gomez-Banuelos et al., Uncoupling interferons and the interferon signature explains clinical and transcriptional subsets in SLE. Cell Rep Med 5, 101569 (2024).

65. J. Tian et al., Dysregulation in keratinocytes drives systemic lupus erythematosus onset. Cell Mol Immunol 22, 83–96 (2025).

66. S. K. Shoffner-Beck et al., Lupus dermal fibroblasts are proinflammatory and exhibit a profibrotic phenotype in scarring skin disease. JCI Insight 9, (2024).

67. S. Braasch, C. Weishaupt, E. Spukti, M. Bohm, S. A. Braun, Scarring Alopecia Under Immune Checkpoint Blockade: a Report of Three Cases. Acta Derm Venereol 102, adv00792 (2022).

68. V. Sibaud et al., Dermatological adverse events with taxane chemotherapy. Eur J Dermatol 26, 427–443 (2016).

69. V. Sibaud, Dermatologic Reactions to Immune Checkpoint Inhibitors : Skin Toxicities and Immunotherapy. Am J Clin Dermatol 19, 345–361 (2018).

70. E. Coleman et al., Inflammatory eruptions associated with immune checkpoint inhibitor therapy: A single-institution retrospective analysis with stratification of reactions by toxicity and implications for management. J Am Acad Dermatol 80, 990–997 (2019).

71. L. Zhou et al., The PD-1/PD-L1 pathway in murine hair cycle transition: a potential anagen phase regulator. Arch Dermatol Res 313, 751–758 (2021).

72. L. Rittie, Cellular mechanisms of skin repair in humans and other mammals. J Cell Commun Signal 10, 103–120 (2016).

73. X. Zhai, M. Gong, Y. Peng, D. Yang, Effects of UV Induced-Photoaging on the Hair Follicle Cycle of C57BL6/J Mice. Clin Cosmet Investig Dermatol 14, 527–539 (2021).

74. A. K. Joseph, L. F. Abbas, B. F. Chong, Treatments for disease damage in cutaneous lupus erythematosus: A narrative review. Dermatol Ther 34, e15034 (2021).

75. A. F. Franca, E. M. de Souza, Histopathology and immunohistochemistry of depigmented lesions in lupus erythematosus. J Cutan Pathol 37, 559–564 (2010).

76. R. J. Garcia et al., Endothelin 3 induces skin pigmentation in a keratin-driven inducible mouse model. J Invest Dermatol 128, 131–142 (2008).

77. S. Tamoutounour et al., Keratinocyte-intrinsic MHCII expression controls microbiota- induced Th1 cell responses. Proc Natl Acad Sci U S A 116, 23643–23652 (2019).

78. L. Fan et al., Antigen presentation by keratinocytes directs autoimmune skin disease. Proc Natl Acad Sci U S A 100, 3386–3391 (2003).

79. Y. Wang et al., A Spatially Coordinated Keratinocyte-Fibroblast Circuit Recruits MMP9(+) Myeloid Cells to Drive IFN-I-Driven Inflammation in Photosensitive Autoimmunity. bioRxiv, (2025).

80. S. Shakiba et al., Spatial characterization of interface dermatitis in cutaneous lupus reveals novel chemokine ligand-receptor pairs that drive disease. bioRxiv, (2025).

81. X. Sun, J. L. Chao, M. Gerner, K. B. Elkon, Induction of Type I Interferon-Dependent Activation and Migration of Inflammatory Dendritic Cells to Local Lymph Nodes by UV Light Exposure. Arthritis Rheumatol 77, 867–875 (2025).

82. G. A. Hile, J. M. Kahlenberg, Immunopathogenesis of skin injury in systemic lupus erythematosus. Curr Opin Rheumatol 33, 173–180 (2021).

83. C. C. Berthier et al., Molecular Profiling of Cutaneous Lupus Lesions Identifies Subgroups Distinct from Clinical Phenotypes. J Clin Med 8, (2019).

84. W. D. Shipman et al., A protective Langerhans cell-keratinocyte axis that is dysfunctional in photosensitivity. Sci Transl Med 10, (2018).

85. Y. Zhao et al., VISTA-induced tumor suppression by a four amino acid intracellular motif. bioRxiv, (2025).

86. T. B. El-Abaseri, L. A. Hansen, EGFR activation and ultraviolet light-induced skin carcinogenesis. J Biomed Biotechnol 2007, 97939 (2007).

87. L. A. Hansen et al., Genetically null mice reveal a central role for epidermal growth factor receptor in the differentiation of the hair follicle and normal hair development. Am J Pathol 150, 1959–1975 (1997).

88. Q. Fu et al., The role of cyclic GMP-AMP synthase and Interferon-I-inducible protein 16 as candidatebiomarkers of systemic lupus erythematosus. Clin Chim Acta 524, 69–77 (2022).

89. T. Li et al., A novel STING1-activating mutation is identified in a patient with childhood-onset systemic lupus erythematosus. Clin Immunol 276, 110493 (2025).

90. W. Xu et al., Immune-Checkpoint Protein VISTA Regulates Antitumor Immunity by Controlling Myeloid Cell-Mediated Inflammation and Immunosuppression. Cancer Immunol Res 7, 1497–1510 (2019).

91. M. H. Kazemi et al., FOXO1 pathway activation by VISTA immune checkpoint restrains pulmonary ILC2 functions. J Clin Invest 135, (2025).

92. Y. Kuchitsu et al., STING signalling is terminated through ESCRT-dependent microautophagy of vesicles originating from recycling endosomes. Nat Cell Biol 25, 453– 466 (2023).

93. C. Q. Lei et al., FoxO1 negatively regulates cellular antiviral response by promoting degradation of IRF3. J Biol Chem 288, 12596–12604 (2013).

94. T. Abe, G. N. Barber, Cytosolic-DNA-mediated, STING-dependent proinflammatory gene induction necessitates canonical NF-kappaB activation through TBK1. J Virol 88, 5328– 5341 (2014).

95. X. Chen, Y. Chen, Ubiquitination of cGAS by TRAF6 regulates anti-DNA viral innate immune responses. Biochem Biophys Res Commun 514, 659–664 (2019).

96. E. C. Nowak et al., The role of VISTA engagement in limiting neutrophil-mediated inflammation. J Immunol, (2025).

97. F. Li, C. A. Adase, L. J. Zhang, Isolation and Culture of Primary Mouse Keratinocytes from Neonatal and Adult Mouse Skin. J Vis Exp, (2017).

